# rcCAE: a convolutional autoencoder method for detecting intra-tumor heterogeneity and single-cell copy number alterations

**DOI:** 10.1101/2022.12.04.519013

**Authors:** Zhenhua Yu, Furui Liu, Fangyuan Shi, Fang Du

## Abstract

Intra-tumor heterogeneity (ITH) is one of the major confounding factors that result in cancer relapse, and deciphering ITH is essential for personalized therapy. Single-cell DNA sequencing (scDNA-seq) now enables profiling of single-cell copy number alterations (CNAs) and thus aids in high-resolution inference of ITH. Here, we introduce an integrated framework called rcCAE, to accurately infer cell subpopulations and single-cell CNAs from scDNA-seq data. A convolutional autoencoder (CAE) is employed in rcCAE to learn latent representation of the cells as well as distill copy number information from noisy read counts data. This unsupervised representation learning via the CAE model makes it convenient to accurately cluster cells over the low-dimensional latent space, and detect single-cell CNAs from enhanced read counts data. Extensive performance evaluations on simulated datasets show rcCAE outperforms existing CNA calling methods, and is highly effective in inferring clonal architecture. Furthermore, evaluations of rcCAE on two real datasets demonstrate it is able to provide more refined clonal structure, of which some details are lost in clonal inference based on integer copy numbers.

## Introduction

### Intra-tumor heterogeneity and single-cell DNA sequencing

The evolutionary history of cancer is often depicted as a phylogenetic tree that elucidates the linage relationship between tumor clones [1, 2, 3, 4, 5, 6]. The emergence of each tumor clone is accompanied with a set of newly acquired genomic mutations such as single-nucleotide variation (SNV) and copy number alteration (CNA), and this genetic intra-tumor heterogeneity (ITH) has been one of the major confounding factors that contribute to therapy resistance and cancer relapse [1, 2, 6]. Without a full understanding of ITH, cancer treatment may often target only major clones whereas the low-prevalence clones may gain a growth advantage after therapy, resulting in cancer relapse [7]. Comprehensively charactering ITH is crucial to understand cancer progression and thus help designing personalized treatment [8, 9, 10]. This requires accurate detection of genomic mutations of each tumor clone as well as inference of the phylogenetic tree.

Single-cell DNA sequencing (scDNA-seq) makes it convenient to profile genomes of single cells and thus enables elucidation of ITH based on single-cell SNVs or CNAs [11]. In scDNA-seq, whole-genome amplification (WGA) is employed to amplify the DNA of isolated single cells to required amount of genetic material for sequencing. WGA tends to introduce amplification bias and errors into the produced sequences, which makes the sequenced reads error-prone and read count fluctuates extensively across genomic regions. The low sequencing coverage and amplification bias make the distribution of read counts over-dispersed, and this poses a critical challenge of how to accurately decipher CNAs from noisy read counts.

### Existing methods for calling CNAs from scDNA-seq data

To date, a few methods have been specifically proposed to detect single-cell CNAs from scDNA-seq data [12, 13, 14, 15, 16, 17, 18, 19, 20]. These methods can be divided into two categories: detect single-cell CNAs with or without normal cells serving as negative controls. Ginkgo [12] is the first CNA calling method specifically designed for scDNA-seq data. It provides a web platform to detect single-cell CNAs using a circular binary segmentation (CBS) [21] algorithm, and also constructs a tumor phylogenetic tree based on the inferred single-cell copy-number profiles. SCNV [13] first identifies normal cells and pools them together as a composite control, then employs a bin-free segmentation approach to estimate copy numbers. It requires at least 20 normal cells to generate the control, and may be not applicable to datasets containing few normal cells. Similarly, SCOPE [14] uses a generalized likelihood ratio test to jointly segment all cells, and leverages a modified Bayesian information criterion (BIC) [22] to determine the optimal number of segments. It automatically identifies normal cells based on calculation of the Gini coefficient and uses them to remove outlier cells. SCICoNE [15] is developed to simultaneously infer single-cell CNAs and a CNA mutation tree. One disadvantage of SCICoNE is that it treats breakpoint detection and CNA analysis as two separate procedures, which may disseminate errors made during breakpoint detection to downstream CNA analysis. SCYN [17] relies on SCOPE to prepare and preprocess read counts, thereafter employs a dynamic programming algorithm to find the optimal segmentation results by maximizing a simplified modified BIC. The authors also introduce SeCNV [20] to perform cross-sample segmentation on normalized read counts. Recently, a method called SCONCE [19] is proposed to call single-cell CNAs using a hidden Markov model (HMM), and it requires normal cells as negative controls to reduce false CNA calls.

Despite to the fact that these methods perform well on their respective test datasets, there are still two problems that are not well addressed in these methods: 1) how to accurately detect single-cell CNAs from noisy and extensively fluctuated read counts without normal cells as negative controls; 2) how to directly infer cell subpopulation composition from the raw read counts rather than the inferred integer copy numbers, thus avoid propagation of errors made in CNA analysis to clonal structure inference. These two problems are intertwined with each other, therefore integrated frameworks for simultaneously inferring single-cell CNAs and cell subpopulations from raw read counts are sorely required for scDNA-seq data.

### Proposed method rcCAE

Deep generative models such as variational autoencoder (VAE) [23, 24] have shown great success in learning representations from single-cell sequencing data [25, 26, 27, 28, 29, 30]. The intuition behind these methods is that observed genetic heterogeneity among single cells actually results from some biological processes that are related to tumor evolution and could be modeled as latent variables in the autoencoders [28]. Inspired by this conception, we develop a novel computational method for analyzing singlecell read counts based on a Convolutional AutoEncoder framework (rcCAE) as shown in Figure 1. rcCAE uses a convolutional encoder network to project normalized singlecell read counts into a low-dimensional latent space where the cells are clustered into distinct subpopulations through a Gaussian mixture model (GMM), and leverages a convolutional decoder network to recover the read counts from learned latent representations. This encoding-decoding process could be treated as a distillation process that extracts copy number information from the noisy read counts by attenuating the effects of non-biological confounding factors, thus makes the recovered data being higher in signal-to-noise ratio. We then apply a novel hidden Markov model (HMM) on the recovered data to simultaneously segment the genome and infer absolute copy numbers for each cell. To the best of our knowledge, rcCAE is the first deep learning-based method for calling single-cell CNAs from scDNA-seq data. We comprehensively evaluate rcCAE on both simulated and real datasets to verify its high scalability on different-sized scDNA-seq datasets, and also compare rcCAE to the state-of-the-art (SOTA) CNA detection methods. The results showcase our method can accurately characterize cell subpopulation composition in latent space, and is more accurate than the competitors in calling absolute copy numbers of single cells.

**Fig. 1.**
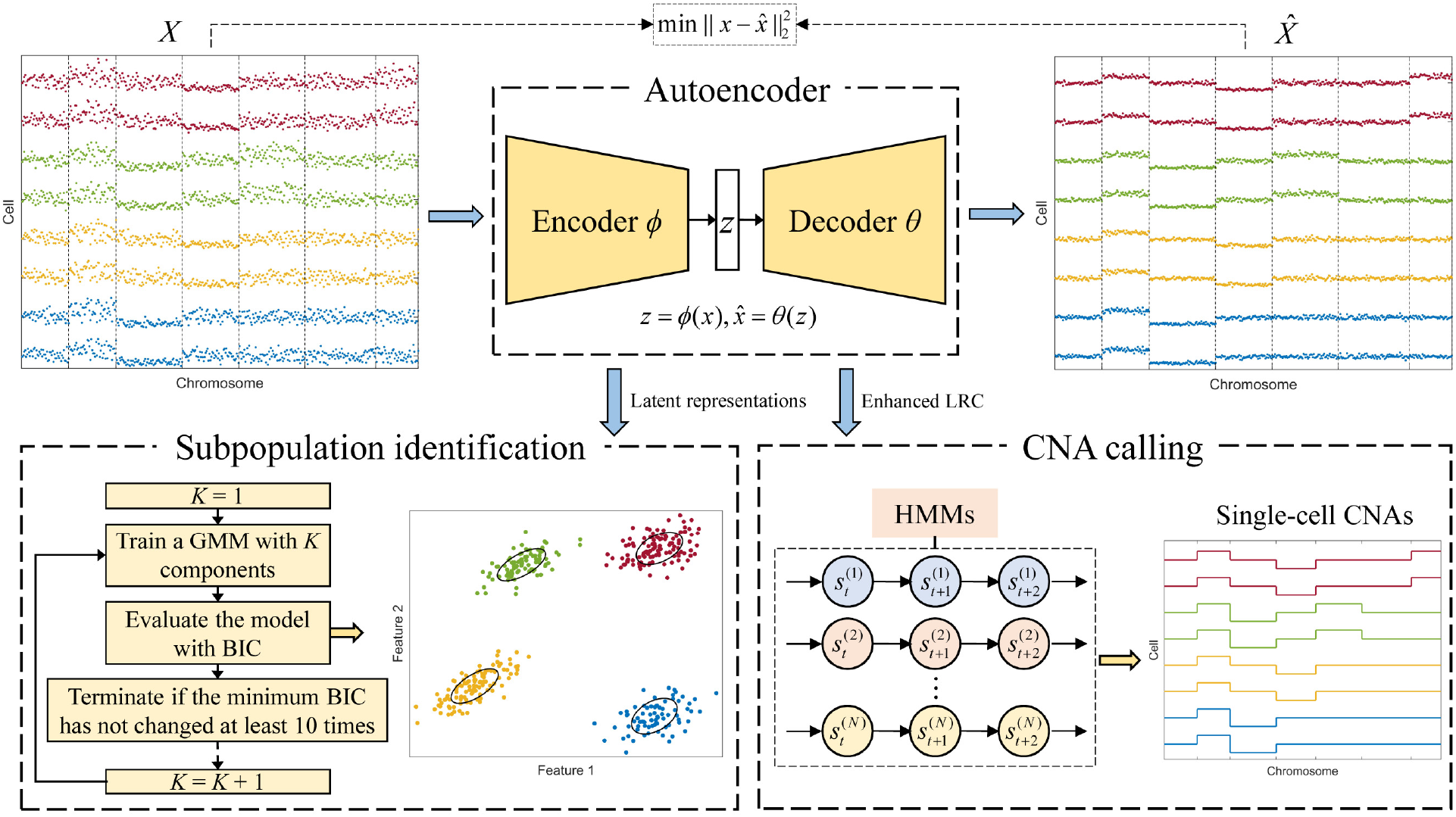
The workflow of rcCAE. rcCAE takes the log2 transformed read counts (LRC) of all cells as inputs, and yields cell subpopulation composition as well as single-cell copy number segments as outputs. With a convolutional autoencoder for unsupervised representation learning, rcCAE projects the LRC data into a latent space where cells are efficiently clustered into distinct subpopulations, and effectively distills copy number information from noisy LRC data. A Gaussian mixture model is employed to identify cell subpopulations based on the latent representation of cells, and the number of subpopulations is determined by Bayesian information criterion. For calling CNAs for each cell, rcCAE employs a hidden Markov model to jointly segment the genome and infer absolute copy number for each segment.

## Results

### The workflow of rcCAE

The framework of rcCAE mainly consists of four components: 1) a module to fetch read counts of fixed-size bins from the BAM files of sequenced cells, and preprocess the read counts; 2) a convolutional autoencoder (CAE) to learn latent representations of single cells and enhance the quality of read counts; 3) a GMM based clustering model to find cell subpopulations; and 4) an HMM to predict copy number segments of each cell. Specifically, encoder of the CAE projects log2 transformed read counts (LRC) of each cell into a lowdimensional space and the decoder recovers the LRC data from the latent space. Given the learned latent features, the GMM model coupled with BIC for model selection is employed to cluster cells into distinct subpopulations. The recovered LRC data from the CAE are then analyzed using the HMM to detect CNAs of each cell. More methodological details of rcCAE are given in Methods section.

### Evaluation on simulated datasets

#### Simulation of scDNA-seq datasets

As human cancers often exhibit aneuploidy, we generate datasets with different tumor ploidy (Simulation A: near-diploidy, Simulation B: near-triploidy and Simulation C: near-tetraploidy). To further investigate potential application of rcCAE to datasets with higher ploidy levels, we also simulate near-pentaploidy datasets (Simulation D). Following the simulation process adopted in [31], we set the minimum size of CNAs to 3Mb, and the maximum size to 20Mb. CNAs are randomly sampled and iteratively inserted into chromosomes 1-3 of human genome hg19 until there are no feasible loci to insert the simulated CNA. We generate 500 cells from 16 subpopulations including a normal subpopulation and 15 tumor clones for Simulations A-C, while generate 100 cells from 5 subpopulations including a normal subpopulation and 4 tumor clones for Simulation D. The simulation is repeated 5 times for each tumor ploidy, and the average numbers of CNAs for Simulations A-D are 18, 35, 36 and 40, respectively. More details about dataset simulation are provided in Supplementary Methods.

#### Performance evaluation

Estimating copy number of a bin can be treated as a classification problem where each copy number is a class. To measure copy number estimation accuracy, we make a comparison between the predicted copy number and the ground truth for each bin, and calculate three metrics for performance evaluation: 1) accuracy: proportion of bins whose copy numbers are accurately predicted; 2) balanced accuracy: mean of per-class accuracies [32]; and 3) macro-averaged F-measure: mean of per-class F1-scores. In addition, we also assess ploidy estimation performance by comparing the predicted average copy number (ACN) with the ground truth value. For inferring tumor clones, we use adjusted rand index (ARI) to measure the clustering accuracy.

When running rcCAE on the simulated datasets, we set bin size to 200kb and the number of epochs for training the CAE to 200 on all simulated datasets as training loss changes little after 200 epochs as shown in Figure S1. Example results of rcCAE on three simulated datasets from Simulations A-C are given in Figures S2-S4. It is observed that rcCAE significantly enhances the quality of LRC by decreasing the variance of LRC in each copy number state. This improvement benefits from the adopted encoding-decoding distillation procedure that makes it convenient to distinguish between different copy number states and between distinct tumor clones.

#### Comparison to other methods

We compare rcCAE to five state-of-the-art CNA calling methods including SCOPE [14], SCYN [17], SCICoNE [15], SeCNV [20] and SCONCE [19] on the simulated datasets from Simulations A-C. All these methods use read counts to call single-cell CNAs and perform well on their respective test datasets. We exclude CHISEL [18] from performance comparison as it uses both read counts and allele frequency data for estimating copy numbers. In addition, we introduce a null model (H0) as the baseline model that predicts the most common copy number (2 for diploid, 3 for triploid and 4 for tetraploid datasets) for all bins. The detailed description about the performance assessment and parameter configuration for the investigated methods are provided in the Supplementary Methods.

We first examine ploidy estimation accuracy of the investigated methods. Difference between the real and predicted ACNs (denoted as ΔACN) is calculated for each cell to indicate the ploidy estimation accuracy. The results in Figure 2 show rcCAE yields better predictions than the SOTAs across different simulations, especially on near-tetraploid datasets. SCICoNE overestimates the ploidy on near-diploid datasets (Simulation A) and predicts all cells to be near-triploid (median ΔACN is −1.02), while other methods perform well with median ΔACN close to 0. On near-triploid datasets (Simulation B), SCOPE, SCYN and SeCNV perform similarly and accurately estimate the ploidy for most of the samples. SCICoNE tends to overestimate the ploidy (median ΔACN is −0.21) and SCONCE erroneously identifies majority of the cells to be near-tetraploid (median ΔACN is 1). By comparison, our method produces accurate inferences of the ploidy for all cells, the mean of ΔACN is 0.0048. On near-tetraploid datasets (Simulation C), all of the existing methods tend to underestimate the ploidy, the median ΔACN of SCOPE, SCYN, SCICoNE, SeCNV and SCONCE are 1.87, 1.87, 0.69, 1.82 and 1.961, respectively, indicating they fail to distinguish between different copy number states on most of the tetraploid samples. Our method is able to assign correct ploidy to each cell, therefore achieves better performance than the competitors. Figure S5 gives a detailed view of the ploidy estimation results of all methods, and the results show rcCAE’ predictions are in concordance with the ground truth.

**Fig. 2.**
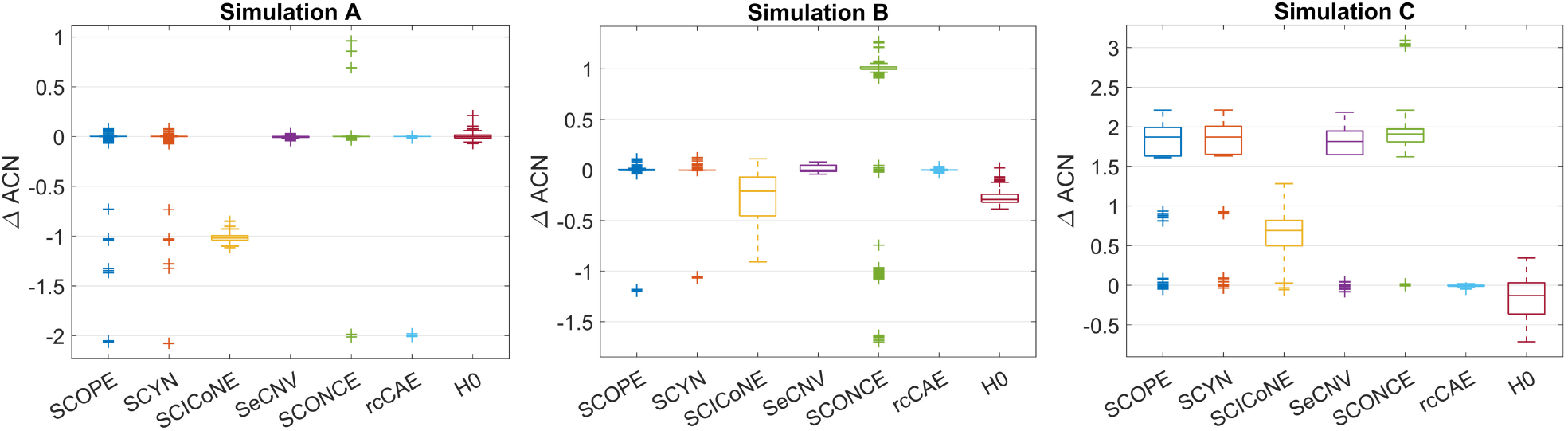
Ploidy estimation accuracy of the investigated methods. Average copy number (ACN) is calculated for each cell, and the difference between real and predicted ACNs (denoted as ΔACN) is used to indicate ploidy estimation accuracy of each method. Results on simulated 5 near-diploid datasets (Simulation A), 5 near-triploid datasets (Simulation B) and 5 near-tetraploid datasets (Simulation C) are analyzed. Each boxplot is generated based on all cells included in the corresponding simulation. The average number of bins used for evaluation is 2509 for Simulation A, 2636 for Simulation B and 2166 for Simulation C. H0 denotes the baseline model that predicts the most common copy number (2 for diploid, 3 for triploid and 4 for tetraploid datasets) for all bins. rcCAE generally performs better than other methods, especially on the near-tetraploid datasets where all of the existing methods tend to underestimate the ploidy.

We also assess how well rcCAE and other methods perform in estimating absolute copy numbers, and Figure 3 shows F-measures of the methods. The copy number accuracy and balanced accuracy are given in Figures S6-7. All performance metrics are calculated per-cell for each method. We use the common bins covered by the results of all methods for evaluation on each dataset. The average number of bins used for evaluation is 2509 for Simulation A, 2636 for Simulation B and 2166 for Simulation C. On near-diploid datasets, SCOPE, SCYN, SeCNV, SCONCE and rcCAE reaches ≥0.993 median F-measure, while SCICoNE fails to identify absolute copy numbers (median F-measure is 0.012) due to overestimation of the ploidy. Similar performance is observed for SCOPE, SCYN, SeCNV and rcCAE on near-triploid datasets. By comparison, SCICoNE yields median F-measure of 0.556 and SCONCE performs poorly as it overestimates the ploidy. On near-tetraploid datasets, the existing methods suffer from degraded performance due to underestimation of the ploidy, and rcCAE surpasses the competitors with median F-measure of 0.935. In addition, we measure the consistency of each method in identifying each copy number state, and the results are shown in Figure S8.

**Fig. 3.**
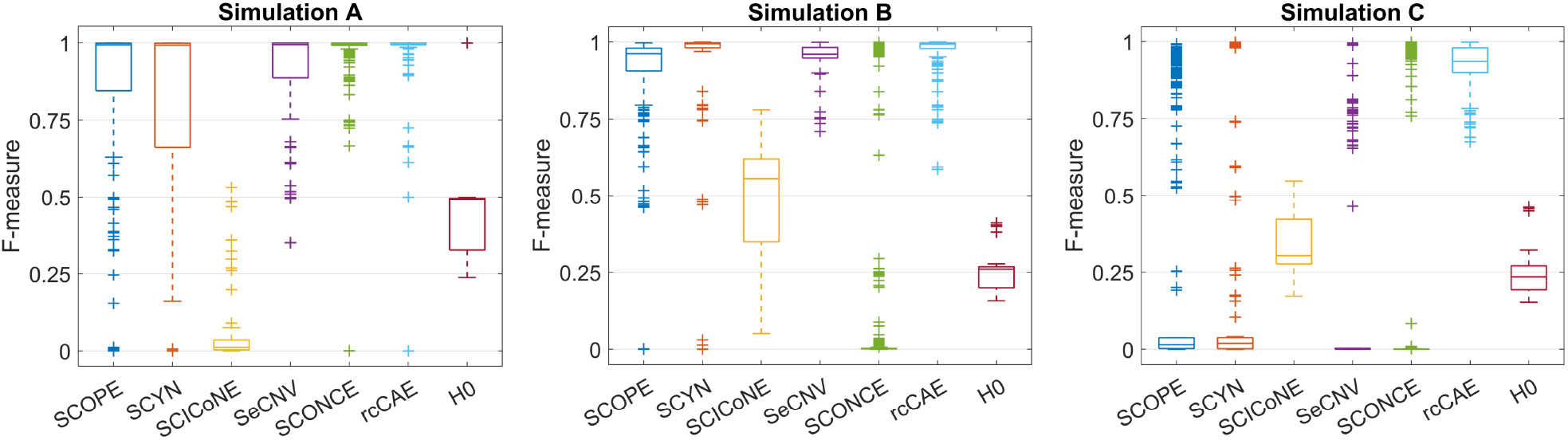
F-measure of the investigated methods. For each cell, the macro-averaged F-measure is calculated as the mean of per-class F1-scores. Results on simulated 5 near-diploid datasets (Simulation A), 5 near-triploid datasets (Simulation B) and 5 near-tetraploid datasets (Simulation C) are analyzed. Each boxplot is generated based on all cells included in the corresponding simulation. The average number of bins used for evaluation is 2509 for Simulation A, 2636 for Simulation B and 2166 for Simulation C.

We proceed to evaluate the performance of rcCAE in identifying tumor clones. Two conventional dimensionality reduction methods including PCA [33] and t-SNE [34] are adopted for comparison by clustering the projected data using the same GMM. More details about how t-SNE is used are provided in the Supplementary Methods. An example of the clustering results is given in Figure S9 that shows rcCAE is more effective in disentangling distinct cell subpopulations. Figure S10 gives ARI scores of all methods, and suggests our method outperforms PCA and t-SNE across different tumor ploidy. rcCAE exactly deciphers clonal composition on 12 out of 15 datasets, and performs acceptedly well on the remaining 3 datasets (ARI > 0.94).

Finally, we assess rcCAE on the simulated near-pentaploidy datasets. An example of copy number estimation results (Figure S11) implies rcCAE significantly improves the quality of LRC data and accurately identifies the most abundant copy number of 5 for all tumor clones. The results in Figure S12 show rcCAE achieves median accuracy of 0.972, balanced accuracy of 0.984 and F-measure of 0.931, which suggests our method is still highly effective in dealing with high-ploidy datasets.

#### The effects of number of cells and hyper-parameters

We assess the effects of number of cells and hyper-parameters on rcCAE’ performance. The results in Figure 4 and Figure S13 suggest our method is able to deliver accurate predictions of copy number (an example is given in Figure S14) and ploidy across different data sizes, and performs acceptedly well in inferring subpopulation composition when the number of cells is larger than 50.

**Fig. 4.**
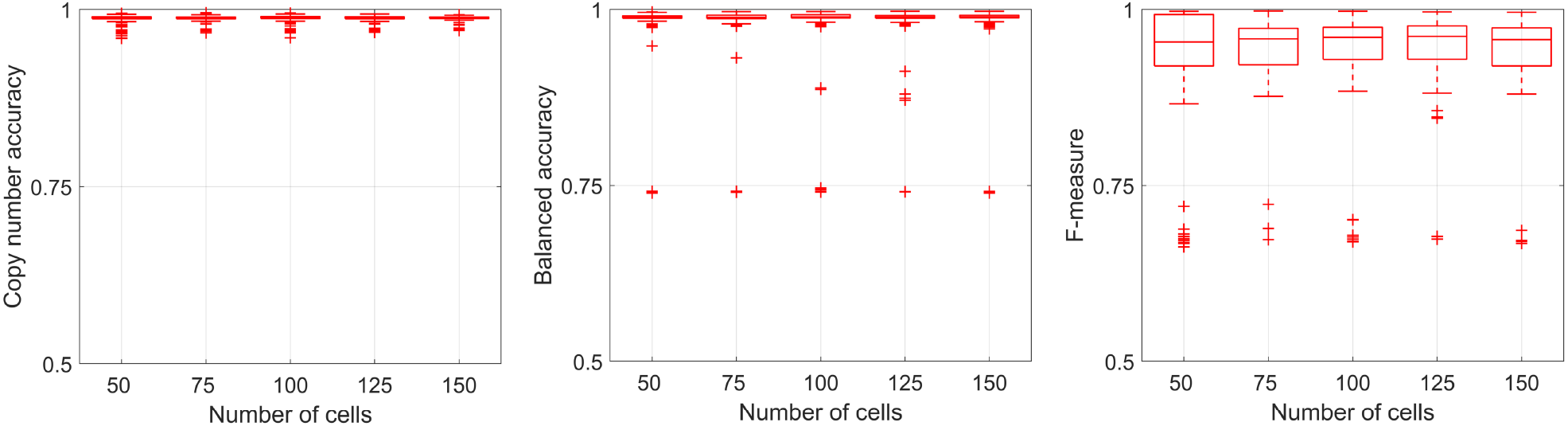
Performance evaluation on different-sized datasets. Values in {50, 75, 100, 125, 150} are tested for the number of cells, and 50 datasets for performance evaluation are generated from a near-diploid dataset in the Simulation A. The cell clustering, copy number as well as ploidy estimation accuracy are measured with respected to the number of cells. Each boxplot is generated based on all cells included in the corresponding datasets. The number of bins used for evaluation is 2955 on all datasets.

We also examine the effects of hyper-parameters including kernel size, latent dimension and bin size on the performance of rcCAE. The results suggest rcCAE reaches consistently good performance across different values of the hyperparameters (Figures S15-20), and detailed description is given in Supplementary Results.

### Evaluation on a breast cancer dataset

We use rcCAE to infer tumor clonal structure from a breast ductal carcinoma dataset consisting of 100 single cells [35]. A previous study [35] has clustered the cells into 4 subpopulations (D+P, H, AA and AB) based on integer copy number profiles, and each subpopulation consists of 47, 24, 25 and 4 cells, respectively. Given the highly different copy number profiles between the subpopulations, we assess if rcCAE could decipher the same clonal structure from single-cell raw read counts data.

Read counts data are obtained for every 500kb bins, and we get a data matrix with size of 100×5120. The number of epochs for training the CAE is set to 200 (as suggested by the results in Figure S21). Our method clusters the cells into 5 subpopulations each with 45, 24, 25, 5 and 1 cells, respectively (Figure 5 and Figures S22-23). Cluster 1 consists of predominantly diploid cells and aligns well with the D+P subpopulation. Cells from cluster 2 are characterized by hemizygous deletions on multiple chromosomes, and the results are in high concordance with the H subpopulation, indicating our method precisely identifies this tumor clone. Cluster 3 is comprised of aneuploid cells that exhibit extensive copy number gains on all chromosomes and hemizygous deletions on chromosomes 4-7. The number of cells is consistent with that of the AA subpopulation, these results demonstrate our method exactly recovers the AA subpopulation. Cluster 4 contains 5 cells of which 4 cells show highly similar copy number profiles, and may be from the AB subpopulation. Our method finds another cluster 5 formed by only one cell that is featured by hemizygous deletions on chromosomes 2, 3q, 8, 9 and 16. This cell is significantly different from the cells of cluster 2 in the copy number profiles (Figure S22), therefore classified into a separate cluster by our method. By directly analyzing the raw read counts data, rcCAE is able to provide more refined clonal structure of which some details are lost in clonal inference based on single-cell integer copy numbers.

**Fig. 5.**
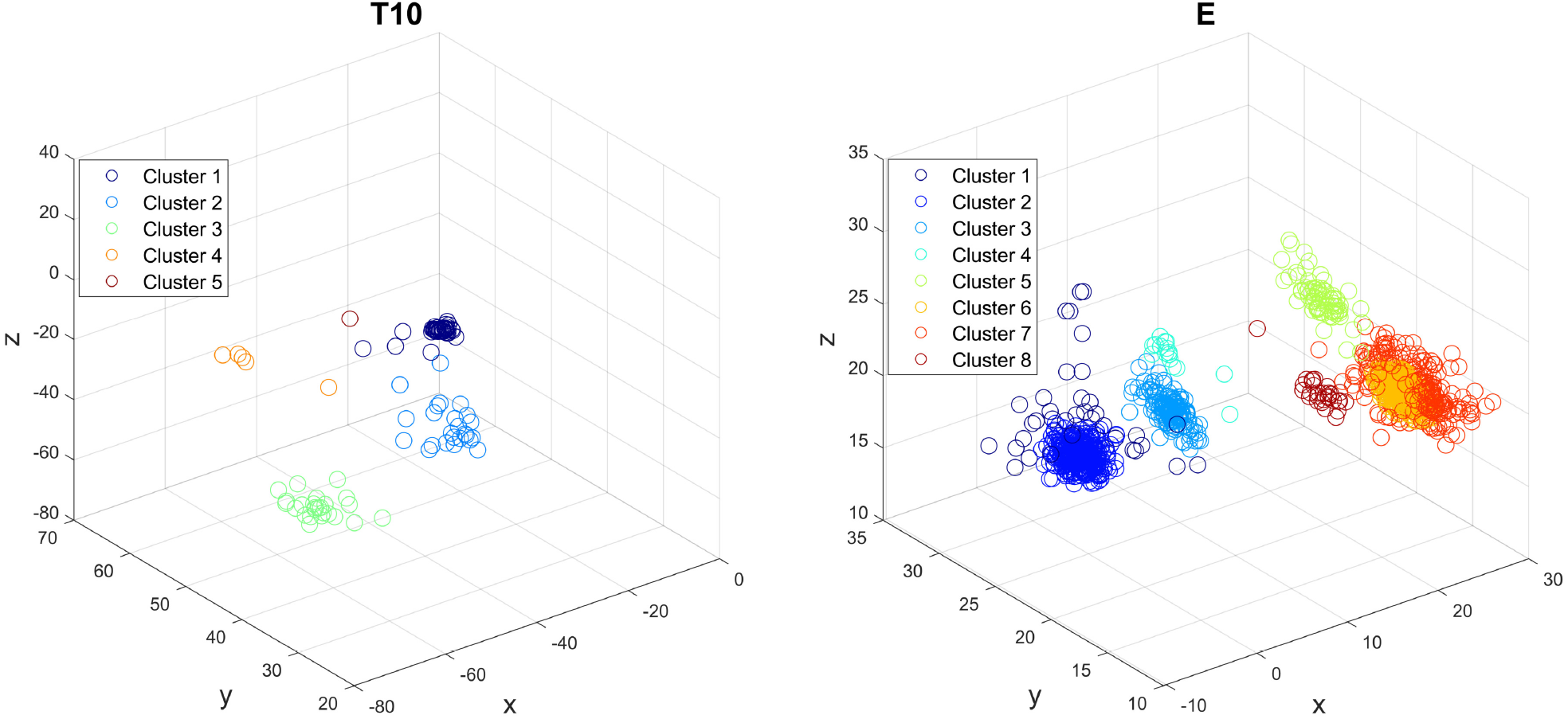
Clustering results of rcCAE on real datasets. On the breast cancer dataset (T10) consisting of 100 cells, rcCAE clusters the cells into 5 subpopulations (labeled as 1-5) each with 45, 24, 25, 5 and 1 cells, respectively (left subfigure). On the 10X Genomics dataset, rcCAE clusters the cells into 8 subpopulations (labeled as 1-8) each with 37, 351, 152, 16, 75, 552, 234 and 29 cells, respectively (right subfigure).

### Evaluation on a 10X Genomics dataset

To further verify the performance of rcCAE on large datasets containing thousands of cells, we apply rcCAE to a 10X Genomics dataset [18]. The cells are obtained from a frozen breast tumor tissue and each cell is sequenced with a coverage between 0.02X and 0.05X. A previous study has assigned 1446/2053 of the cells to 6 clusters including one diploid subpopulation (containing 388 cells) and 5 aneuploid tumor clones (each with 168, 58, 20, 782 and 30 cells, respectively) using CHISEL algorithm [18]. Tumor clones 2 and 3 show highly consistent copy number profiles across the whole genome but different allele-specific copy numbers on chromosome 2, therefore we merge the two clones into a same cluster as they are indistinguishable from each other solely based on the total copy numbers. We now label the new clusters with I-V (each with 388, 168, 78, 782 and 30 cells, respectively). We employ rcCAE to infer clonal structure from this dataset and check if our method could yield similar results as CHISEL by only using read counts data.

The bin size is set to 500kb, and the number of epochs for training the CAE is set to 50 (as shown in Figure S24). rcCAE assigns the cells into 8 clusters (labeled as 1-8) each with 37, 351, 152, 16, 75, 552, 234 and 29 cells, respectively (Figure 5 and Figure S23). A comparison between the original and enhanced LRC data is given in Figure 6 and the results show our method significantly improves the data quality. We find clusters 1 and 2 are diploid subpopulations having highly similar copy number profiles and the total number of cells from them is equal to that of CHISEL cluster I, which suggests rcCAE subdivides the cells of CHISEL cluster I into two sets that are in close proximity in the latent space (Figure 5). Similar results are observed for cluster 3 and 4 (labeled as clone A) exhibiting consistent copy numbers and aligning well with the CHISEL cluster II. Cluster 5 (labeled as clone B) has copy number of 4 on chromosomes 2-3 and copy number of 3 on chromosome 8, and presents very similar copy number profiles with the CHISEL cluster III. The results indicate rcCAE effectively deciphers this clone by identifying 75/78 of the cells. Clusters 6 and 7 (labeled as clone C) also show quite similar copy number profiles and have copy number of 3 on chromosome 2 when compared to clone B. In addition, 786 cells in clone C completely encompasses the 782 cells from the CHISEL cluster IV, which suggests clone C aligns well with the CHISEL cluster IV. Cluster 8 represents a minor clone (labeled as clone D) that contains 29 cells, and we found clone D is in concordance with the CHISEL cluster V. Taken together, rcCAE may be sensitive to LRC changes among the cells belonging to same cluster and tends to generate over-segmented clustering results, however it is still able to accurately recover the copy number profiles of single cells by exploiting strong reconstruction ability of the CAE model.

**Fig. 6.**
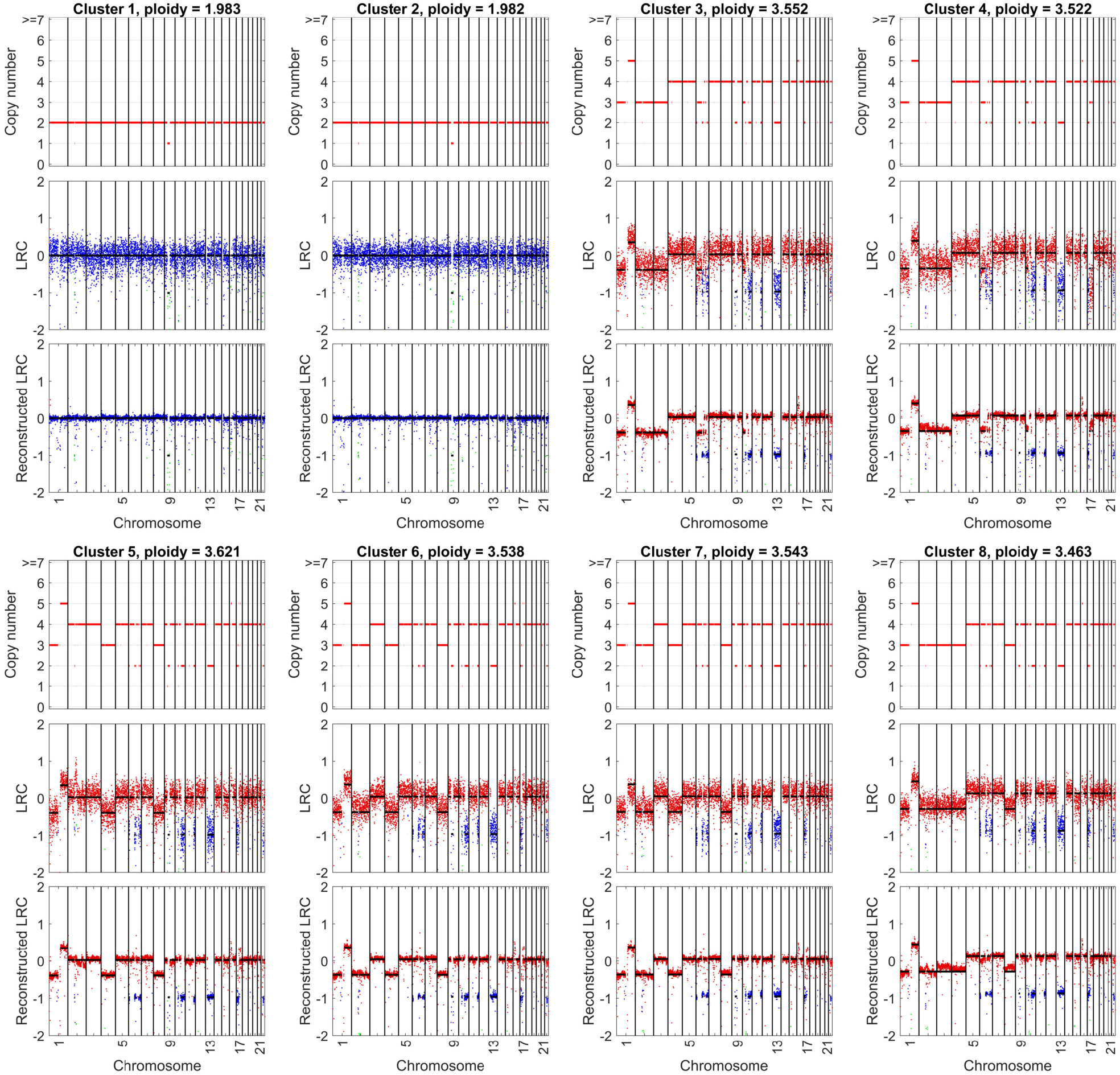
Copy number estimation results of rcCAE on the 10X Genomics dataset. Clusters 1 and 2 are diploid subpopulations having highly similar copy number profiles. Clusters 3 and 4 showcase consistent copy numbers with average ploidy of 3.55 and may come from a same clone (labeled as clone A). Cluster 5 represents another clone (labeled as clone B) that has copy number of 4 on chromosomes 2-3 and copy number of 3 on chromosome 8. Clusters 6 and 7 show quite similar copy number profiles, and may derive from a same clone (labeled as clone C). Compared to clusters 1-7, cluster 8 represents a minor clone (labeled as clone D), and it has copy number of 3 on chromosomes 2-3. The quality of LCR data is significantly enhanced by rcCAE’s encoding-decoding framework, thus single-cell CNAs could be accurately detected.

## Discussion and conclusions

In this paper, we develop rcCAE, an autoencoder based method for jointly inferring ITH and single-cell copy numbers from scDNA-seq data. The approach leverages a convolutional autoencoder to learn latent representations of single cells as well as enhance the quality of read counts, thus provides an integrated framework that enables efficient characterization of underlying cell subpopulations in a low-dimensional latent space, and accurate detection of single-cell copy numbers from improved read counts. Compared to ITH inference based on integer single-cell copy numbers, rcCAE directly deciphers ITH from original read counts, which avoids potential error propagation from copy number analysis to ITH inference. For estimating copy number segments for each cell, rcCAE uses a novel hidden Markov model to explicitly represents tumor ploidy as a key parameter in the emission models to find the solution that best explains the observed read counts. rcCAE is the first method that borrows both the strong representation learning ability of autoencoder and good explanation capability of conventional statistical methods for analyzing single-cell read counts, thus opens a new paradigm for CNA-based ITH analysis.

Further improvements in single-cell CNA calling could be achieved by cross-cell segmentation of the read counts. First, cells in same clone have a common set of breakpoints, and incorporating this information into CNA detection could aid in better identifying breakpoints. Second, cells from different clones may share same breakpoints since tumor evolves by accumulating mutations [36], and cross-cell segmentation could attenuate the effects of read counts fluctuation and thus deliver better segmentation results. A more challenging task is to perform cross-cell segmentation along with identification of absolute copy numbers. Existing methods that are not based on HMMs often require a post-processing procedure to estimate absolute copy numbers after obtaining segmentation results. Despite to the fact that HMM-based methods can provide segmentation as well as absolute copy numbers for a single cell, how to apply HMM to multiple cells is still an open question, and a major challenge is that hidden space size may grow exponentially with the number of cells. Taken together, cross-cell segmentation with simultaneous identification of absolute copy numbers is a promising direction for future study of ITH inference from scDNA-seq data.

## Methods

### Obtain and preprocess read counts

We divide the genome into fix-sized bins and obtain read count of a bin by counting the reads mapped to the bin. BamTools [37] is used to fetch information of each read alignment. We also calculate GC percent and mappability score of each bin to correct read counts bias as well as exclude outlier bins. Specifically, bins that have mappability score of ≥0.9 or GC percent of ≤10% are excluded from downstream analysis [38]. To make read counts comparable among cells, we calculate mean of read counts across all bins for each cell, and divide read count of each bin with the mean. We further exclude bins with mean value of the normalized read counts being in the lower or upper 1% quantile. Considering cells may end up with low sequencing coverage or high amplification bias, we use the normalized read counts to calculate the Gini coefficient for each cell, and eliminate cells having ≥0.3 Gini coefficient.

Finally, per-cell correction for GC-content bias using a median normalization approach [39] is conducted to improve the quality of read counts. For computational convenience, we calculate the log2 transformed read counts of all cells for downstream analysis, here denoted by a *N* × *M* matrix *X* (*N* denotes the number of cells and M is the number of bins).

### Learn latent representations and enhance the quality of LRC

Inspired by recent success of deep generative models in learning representations of cells from single-cell RNA sequencing (scRNA-seq) data [26, 27], we introduce a convolutional autoencoder based approach to represent single cells in a latent space where cells could be efficiently clustered into distinct subpopulations. Formally, for each *x* (one row of the *X*), we use the encoder *ϕ* of the CAE model to get its latent representation *z* = *ϕ*(*x*), and recover *x* through the decoder network *θ* of the CAE, i.e. *x* = *θ*(*z*). Here the latent *z* could be assumed to indicate underlying biological processes that result in the observed cell and are related to mutational events shared by other cells derived from the same tumor clone. The data reconstruction could be treated as a refinement process that distills copy number information from original LRC data, therefore make it convenient to accurately estimate copy numbers from the recovered LRC.

We train the CAE model to minimize the reconstruction loss 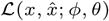 which is measured using the *ℓ*_2_-norm:

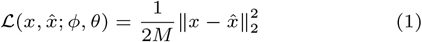

During each training iteration, the mean loss is calculated for a batch of samples, and conventional gradient decent algorithm is employed to update the network parameters. After the model training is completed, the latent representations and recovered LRC data of all cells are inferred as *Z* and 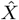, respectively.

### Infer cell subpopulations

Given the latent representations *Z* of all cells, we employ a GMM based clustering approach to find cell subpopulations by following a previous study [28] that uses a GMM to cluster cells based on scRNA-seq data. Specifically, the probability density function of *z* is defined as follows:

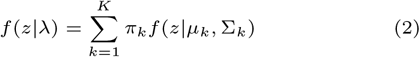

where *K* is the number of clusters, *π_k_* is the proportion of the *k*-th cluster, *f* (*z*|*μ_k_*, ∑_*k*_) is the density function associated with the *k*-th multivariate Gaussian distribution with mean *μ_k_* and covariance matrix ∑_K_, and *λ* = (*π*_1_, *μ*_1_, ∑_1_,…, *π_K_, μ_K_, ∑_K_*) are model parameters.

We adopt expectation maximization (EM) algorithm to update the model parameters, and assign each cell to the cluster having the maximum posterior probability. To determine the best number of clusters (the value of *K*), we calculate the BICs of the models associated with different values of *K*, and select the model having the minimum BIC as the best solution. For implementation, the value of *K* is initialized to 1 and then iteratively increased by one until the minimum BIC has remained unchanged at least 10 times. After the best model is found, the cells can be divided into distinct subpopulations, and cells belonging to the same subpopulation are considered to have highly similar copy number profiles across whole genome.

### Detect single-cell CNAs

To jointly segment the genome and infer absolute copy numbers for each cell, we employ an HMM to analyze the enhanced LRC data. The HMM defines 11 hidden states each of which represents a copy number that ranges from 0 to 10. Here we use 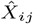 to denote the LRC of the *i*-th cell in the *j*-th bin. We assume 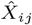 follows a normal distribution under hidden state *s*:

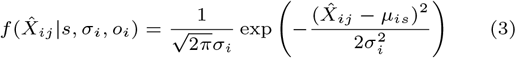

with the mean *μ_is_* defined as:

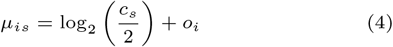

where *o_i_* denotes baseline shift of the LRC due to change of ploidy, *σ_i_* is a global parameter representing the standard variance of LRC, and *c_s_* represents copy number in state *s*. The parameter *o_i_* is introduced to recognize the most abundant copy number state. The value of *o* should be close to 0 for diploid samples, and lower than 0 for hyperploid samples. We use EM algorithm to estimate the model parameters. The expected partial log-likelihood is formulated as:

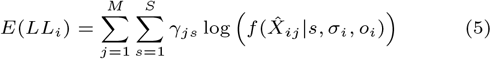

where *γ_js_* is the posterior probability that the *j*-th bin is in state *s*, and *S* denotes the number of hidden states. During the *n*-th iteration of the EM algorithm, we update parameter *o_i_* with the following formula:

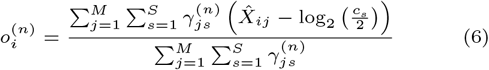

and update *σ_i_* by:

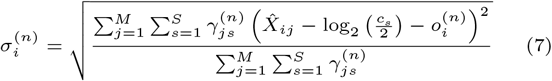

We test different initial values for *o_i_* to check which ploidy should be the most abundant copy number, and initialize *σ_i_* to the standard variance of LRC of the *i*-th cell. After the algorithm converges, the copy number of each bin is deduced from the state that has the maximum posterior probability. We also propose a post-processing step to recalibrate the copy number of highly amplified regions, and more details about the implementation are provided in the Supplementary Methods.

#### Key Points

- We develop a convolutional autoencoder based method rcCAE, which jointly detects cell subpopulations and single-cell CNAs from raw read counts.
- rcCAE exploits the encoding-decoding process to distill copy number information from extensively fluctuated read counts.
- rcCAE is able to directly infer cell subpopulations from read counts, and thus could give more robust characterization of ITH.

## Authors’ contributions

Z.Y. and F.D. conceived the study. Z.Y. developed the algorithm and drafted the manuscript. F.L. and F.S. analyzed the data. All authors read and approved the final manuscript for publication.

## Ethics approval and consent to participate

Not applicable.

## Software availability

The source code and detailed documentation of rcCAE are available at https://github.com/zhyu-lab/rccae.

## Data availability

The breast cancer dataset is available from NCBI SRA with accession number SRA018951, and the 10X Genomics dataset can be freely downloaded from https://support.10xgenomics.com/si_cell-dna/datasets/1.0.0/breast_tissue_E_2k.

**Zhenhua Yu** is an associate professor at School of Information Engineering, Ningxia University. His research interests include bioinformatics, computational biology and deep learning.

**Furui Liu** is a Master’s student at School of Information Engineering, Ningxia University. His research interests include bioinformatics and cancer genomics.

**Fangyuan Shi** is an assistant professor at School of Information Engineering, Ningxia University. Her research interests include bioinformatics and computational biology.

**Fang Du** is a professor at School of Information Engineering, Ningxia University, Yinchuan, China. Her research interests include biomedical big data analysis and knowledge graph.

