## Supplementary Material for "rcCAE: a convolutional autoencoder method for detecting intra-tumor heterogeneity and single-cell copy number alterations"

Zhenhua Yu[1,2,*], Furui Liu[1], Fangyuan Shi[1,2] and Fang Du[1,2]

[1]School of Information Engineering, [2]Collaborative Innovation Center for Ningxia Big Data and Artificial Intelligence, Ningxia University, Yinchuan 750021, China

*To whom correspondence should be addressed.

Contact:

### 1. Supplementary Methods

#### 1.1. Simulate single-cell DNA sequencing data

Simulation of single-cell DNA sequencing (scDNA-seq) data consists of three steps: (1) mimic a clonal tree that depicts lineage relationship between tumor clones; (2) construct genome sequence of each cell given simulated CNAs; (3) generate sequencing reads of each cell according to parameters such as sequencing coverage and read length. Each node except root of the clonal tree represents a tumor clone, and each edge is labeled with new CNAs acquired by the child node. The clonal tree is constructed as follows: (1) attach the first clone to root, and iteratively attach remaining clones to randomly selected non-root nodes of the tree; (2) generate CNAs (size of the CNAs ranges from 3Mb to 20Mb) and assign them to edges of the tree, the number of CNAs assigned to each edge is proportional to weight of the edge (the edge that connects root and the first clone is given a higher weight of 3, and others are given the same weight of 1). To emulate different tumor ploidy $p$, we give higher weight to copy number $p$ when generating CNAs. We first assign 60% out of the CNAs having copy number of $p$ to the first edge of the tree, then process remaining CNAs according to the weights of the edges; (3) uniformly attach 20% of all cells to the nodes of the tree, and iteratively add remaining cells to the tree. The probability of selecting a node as the attachment point is proportional to current size of the node, which is defined as the number of cells already added to the node, and this promises generation of different-sized subpopulations including major and minor clones. Given the simulated clonal tree, the CNAs of each cell can be obtained by accumulating the mutations along the path from root to the cell. We employ SCSsim [1] software to construct the genome sequence and generate sequencing reads (sequencing coverage is set to 0.03) of each cell. FASTQ files are processed using BWA [2] with default parameters to generate read alignments, and BAM files are generated using SAMtools [3].

Datasets with different tumor ploidy (Simulation A: near-diploidy, Simulation B: near-triploidy, Simulation C: near-tetraploidy and Simulation D: near- pentaploidy) are generated. The number of tumor clones is set to 15 and the number of cells is set to 500 for Simulations A-C, while 4 tumor clones and 100 cells for Simulation D. For each simulation, 5 datasets are produced for evaluation.

#### 1.2. Estimate copy numbers of highly amplified regions

rcCAE employs an HMM that considers the maximum copy number of 10 to segment the genome and estimate copy number of each segment, while some highly amplified regions may have copy number of >10. To assign correct copy numbers to these regions, we employ a post-processing approach to recalculate copy number for each segment based on the learned model parameters. Formally, the copy number of the $k$-th segment of the $i$-th cell is tuned as $c_{ik} = round(2^{m_{ik}-o_i+1})$, here $m_{ik}$ denotes the median LRC of the segment, $o_i$ is the estimated baseline shift of LRC and $round(x)$ takes the nearest integer of the input $x$.

#### 1.3. Implementation

For calling single-cell CNAs, we get read counts from scDNA-seq data, and calculate GC-content & mappability from reference genome sequence. This data preparation process provides required input data for downstream analysis.

We use a convolutional autoencoder (CAE) model to learn low-dimensional latent representations of cells and improve the quality of LRC data. The network structure of the CAE is defined as follows: the encoder consists of three 1-D convolutional layers each with 128, 64 and 32 kernels, respectively, the latent layer is a 3-dimensional fully connected layer, and the decoder consists of a fully connected layer and three transposed 1-D convolutional layers each with 64, 128 and 1 kernels, respectively. Leaky Rectified Linear unit (LeakyReLU) activation function is used in all convolutional layers except the last layer of the decoder, and kernel size is set to 7. We use 'Adam' algorithm to train the network with learning rate of 0.0001, and set batch size to 64. The number of epochs is specified per dataset.

Based on the learned latent representations, cells are clustered into distinct subpopulations using a Gaussian mixture model. To determine the best number of clusters (the value of $K$), we calculate the BICs of the models associated with different values of $K$, and select the model having the minimum BIC as the best solution. Specifically, the value of $K$ is initialized to 1 and then iteratively increased by one until the minimum BIC has remained unchanged at least 10 times.

Given the LRC data of a cell, we used an HMM to segment the bins as well as estimate absolute copy numbers of all segments. To find the ploidy that best explains the observed LRC, we introduce a parameter $o$ to model the baseline shift of LRC due to change of ploidy. The value of $o$ should be close to 0 for diploid samples, and lower than 0 for hyperploid samples, therefore we test different initial values for $o$ to find the most abundant copy number.

The data preparation module is implemented in C++, the CAE model as well as cell clustering method are implemented in Python, and the HMM is implemented in MATLAB. All experiments are conducted on a server with 1 RTX 2080 Ti GPU, 128GB RAM and 64 CPU cores.

#### 1.4. Details of investigated methods

##### 1.4.1. SCOPE

SCOPE [4] uses a generalized likelihood ratio test to jointly segment all cells, and is implemented in R. When running SCOPE on simulated datasets, we set parameter "autochr" to 3 in "get_bam_bed.R" file as our simulated data only contain chromosomes 1-3, and specify parameter "resolution" to 200 in "run_scope.R" file, while set other parameters to their default values. The SCOPE software is available at https://github.com/rujinwang/SCOPE.

##### 1.4.2. SCYN

SCYN [5] employs a dynamic programming algorithm to estimate copy number segments by maximizing a simplified mBIC, and uses SCOPE to prepare and preprocess the read counts. It is also implemented in R. When running SCYN on simulated datasets, we adopt parameter configuration as "--seq paired-end --bin_len 200 --ref hg19 --reg .*[0-9].bam$ --mapq 40". The SCYN software is available at https://github.com/xikanfeng2/SCYN.

##### 1.4.3. SCICoNE

SCICoNE [6] is able to simultaneously infer single-cell CNAs and a CNA mutation tree. It first detects breakpoints and then calls CNAs as well as a CNA tree. SCICoNE is implemented in C++ and Python. When running SCICoNE on simulated datasets, we first generate a bed file defining 200kb bins, and obtain read counts using BedTools [7]. We then execute "breakpoint_detection", "segment_counts" and "inference" commands by specifying "--n_bins" and "--n_cells" parameters while using default parameters for other parameters. The SCICoNE software is available at https://github.com/cbg-ethz/SCICoNE.

##### 1.4.4. SeCNV

SeCNV [8] is a method to perform cross-sample segmentation on single-cell read counts. It is implemented in Python. We use parameter configuration "-r hg19 -b 200000 -p *[0-9].bam" when running SeCNV on simulated datasets. The SeCNV software is available at https://github.com/deepomicslab/SeCNV.

##### 1.4.5. SCONCE

SCONCE [9] employs a hidden Markov model (HMM) to detect CNAs for tumor cells. It is implemented in C++ and R. When running SCONCE on simulated datasets, we first generate a bed file defining 200kb bins, and obtain read counts using BedTools [7]. We then execute "avgDiploid.R", "fitMeanVarRlnshp.R" and "sconce" command under the default parameters. The SCONCE software is available at https://github.com/NielsenBerkeleyLab/sconce.

##### 1.4.6. t-SNE

For running t-SNE [10] on simulated datasets, we first use PCA [11] to reduce the dimension of LRC data to 50, then employ t-SNE to project the reduced data into desired latent dimension. For each simulated dataset, the "learning_rate" parameter is set to 100 and "perplexity" parameter in {10, 20, 30, 40, 50} are tested for t-SNE to select the solution that yields the highest ARI score.

#### 1.5. Usage of rcCAE

The pipeline of rcCAE consists of three steps: 1) fetch single-cell read counts from scDNA-seq data and get GC-content & mappability from reference genome sequence; 2) learn latent representations of cells, obtain quality-enhanced log read counts data, and cluster cells into distinct subpopulations; and 3) call single-cell CNAs from LRC data. In the following sections, we briefly explain the main aspects of the software, and more details about how to install and use rcCAE can be found at https://github.com/zhyu-lab/rccae.

##### 1.5.1. Get read counts, GC-content and mappability

rcCAE fetches single-cell read counts of fixed-size bins from a merged BAM (barcoded BAM as done in 10X Genomics) file or per-cell BAM files. In a barcoded BAM file, reads must be labelled by a barcode, and rcCAE assumes the barcodes are defined according to the 10X Genomics format. In per-cell BAM files, read alignments of each cell are stored in a separate BAM file. rcCAE also calculates GC-content and mappability from reference genome sequence to filter and improve read counts. We implement a tool called "prep" using C++ to prepare all these required data.
The inputs to the tool include: 1) A merged BAM file (10X Genomics) containing sequencing data of all cells or separate BAM files of single cells; 2) A FASTA file defining reference genome sequence; 3) A BIGWIG file for calculating mappability scores; and 4) A barcode file listing the barcodes of all cells (for 10X Genomics) or names of all BAM files to be analyzed (for per-cell BAMs). The reference sequence file formatted as .fasta can be downloaded from UCSC genome browser (http://hgdownload.soe.ucsc.edu/goldenPath/hg19/bigZips/ for hg19, https://hgdownload.cse.ucsc.edu/goldenpath/hg38/bigZips/ for hg38). BIGWIG files formatted as .bw for human genomes (e.g. wgEncodeCrgMapabilityAlign36mer.bw.gz) are also available from UCSC genome browser (https://hgdownload.soe.ucsc.edu/goldenPath/hg18/encodeDCC/wgEncodeMapability/ for

hg18, and https://hgdownload.soe.ucsc.edu/goldenPath/hg19/encodeDCC/wgEncodeMapability/ for hg19), and the BIGWIG files can also be created by following our provided guidelines at rcCAE homepage. For a barcoded BAM, the format of barcode file looks like as follows:

*AAACCTGCACCAAAGG-1*

*AAACCTGCAGGACCAA-1*

*AAACCTGGTAACTTCG-1*

*AAACCTGGTACCAGTT-1*

Each row represents the barcode of a cell. For per-cell BAM files (e.g. cell1.bam, cell2.bam,…), the format of barcode file looks like as follows:

*cell1*

*cell2*

*cell3*

*cell4*

Each row denotes the name (without suffix) of a BAM file. The output of the tool is a single file containing read counts, GC-content and mappability data.

The command to run the tool follows the form:

*./prep/bin/prepInput -b /path/to/sample.bam -r /path/to/hg19.fasta -m /path/to/hg19.bw -B /path/to/barcodes.txt -o example.txt*

##### 1.5.2. Learn latent representations of cells and cluster cells into subpopulations

rcCAE employs a convolutional autoencoder (CAE) model to learn latent representations of cells as well as improve the quality of LRC data. Based on the latent representations, a Gaussian Mixture Model (GMM) based clustering method is used to identify cell subpopulations. The input to the CAE model is the output file of "prepInput" command as described above. The outputs of the CAE include: 1) a file named "latent.txt" to save the learned latent representations of cells; 2) a file named "label.txt" to save the clustering results; 3) a file named "lrc.txt" to store the quality improved LRC data; and 4) a file named "loss.txt" to save the training loss at each epoch. The "lrc.txt" file will be used to call CNAs for each cell.

We implement the CAE model using Python, and the command to train the model follows the form:

*python ./cae/train.py --input ./data/example.txt --epochs 200 --batch_size 64 --lr 0.0001 --latent_dim 3 --seed 0 --output data*

##### 1.5.3. Detect single-cell CNAs

In rcCAE, a hidden Markov model (HMM) is employed to call per-cell CNAs. The inputs to the HMM includes: 1) the "lrc.txt" file outputted by the CAE model; 2) a directory to save the results; and 3) the maximum copy number to consider in the HMM. We implement the HMM using MATLAB, and run the program like the following form:

*SCHMM('lrc.txt','results',10)*

Finally, we also provide a script "run_rccae.sh" to integrate above three steps to run rcCAE, which makes it convenient for users to analyze tumor scDNA-seq data.

### 2. Supplementary Results

#### 2.1 The effects of number of cells

To check the robustness of rcCAE' performance against data size, we apply it to simulated datasets where the number of cells is in {50, 75, 100, 125, 150}. The datasets are generated from a near-diploid dataset in Simulation A. Formally, we first randomly select 5 out of 16 subpopulations, then sample cells from each selected subpopulation with the probability proportional to subpopulation size. This process is repeated 10 times to construct a total of 50 datasets for evaluation. The results suggest rcCAE performs acceptedly well in inferring subpopulation composition when the number of cells is larger than 50, therefore we recommend using rcCAE on datasets containing >50 cells to obtain reliable prediction results.

#### 2.2 The effects of hyper-parameters

The size of convolutional kernels and dimension of latent space are two important hyper-parameters defining the structure of the CAE. To examine the effects of kernel size (denoted as $k$) and latent dimension (denoted as $d$) on the performance of rcCAE, we compare cell clustering and CNA calling results for different configurations of $k$ and $d$. Specifically, values in {3, 5, 7, 9, 11} and {2, 3, 4, 5, 6} are tested for $k$ and $d$, respectively. Copy number estimation results (Figure S15) imply rcCAE performs consistently well in identifying single-cell CNAs regardless of different network structures defined by $k$ and $d$. This high robustness to network structures benefits from rcCAE's effective distillation of copy number information from noisy LRC data and accurate ploidy estimations (Figure S16). By comparison, the hyper-parameters $k$ and $d$ have significant effects on clustering results (Figure S17). Generally, setting latent dimension to 3 or 4 gives more accurate identifications of cell subpopulations, and median-sized kernels tend to deliver more stable clustering results. To make a good tradeoff between inference accuracy and efficiency, we therefore set $k$ to 7 and $d$ to 3 on all experiments.

We also examine the effects of bin size on the inference results of rcCAE. Specifically, values in {100kb, 200kb, 500kb} are tested for bin size. The results suggest rcCAE' performance is robust to the change of bin size, and yields accurate estimation of copy numbers, ploidy and subpopulations (Figure S18-20).

### 3. Supplementary Figures

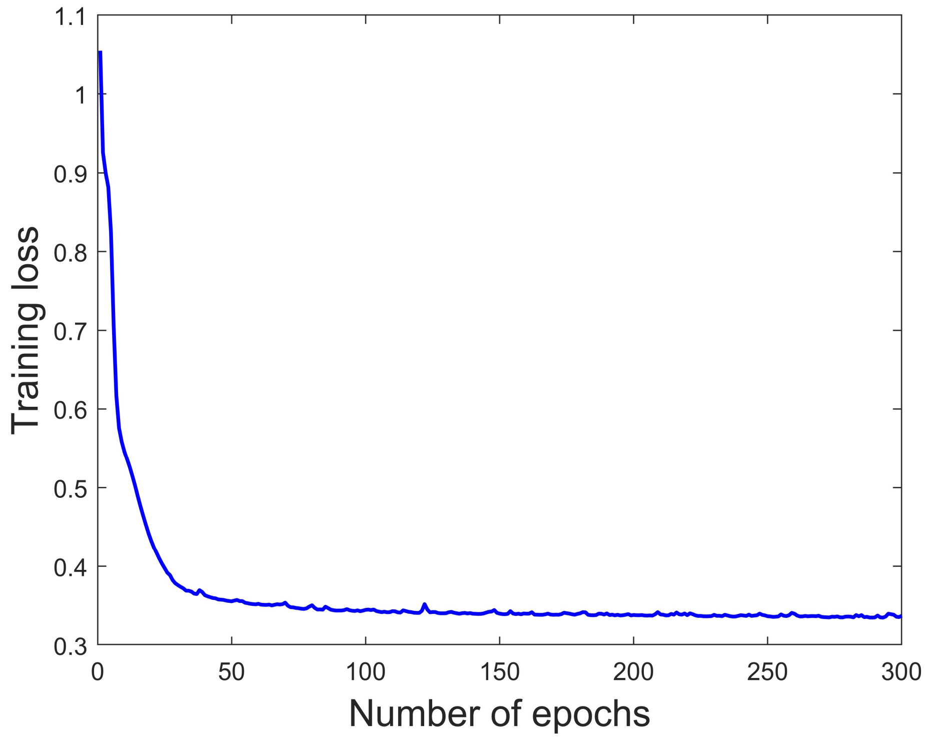

**Figure S1.** Training loss with respect to the number of epochs on a simulated dataset. It is observed that training loss changes litter after 200 epochs. Similar results are observed on other simulated datasets, therefore the number of epochs for training the convolutional autoencoder (CAE) is set to 200 on all simulated datasets.

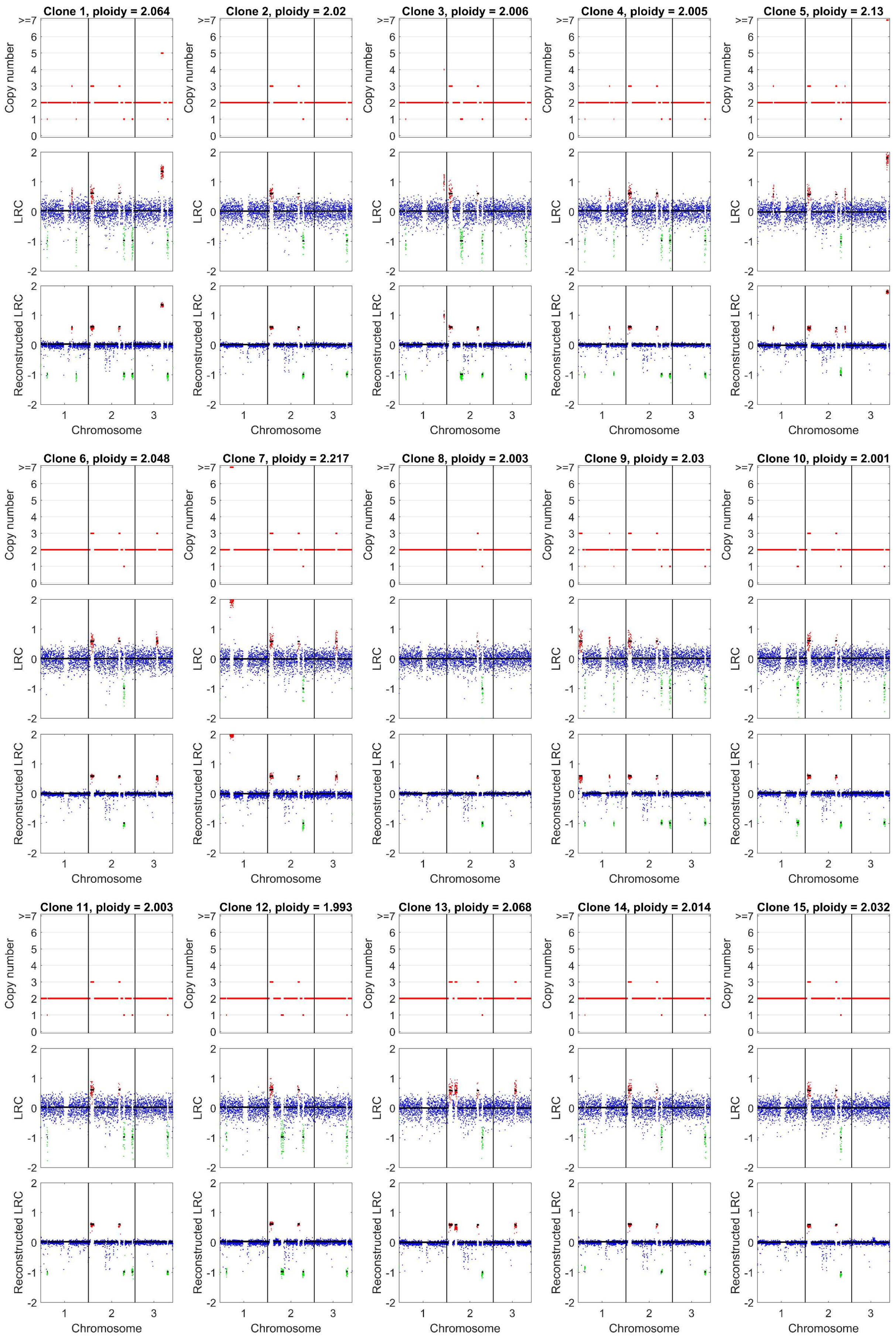

**Figure S2.** Copy number estimation results on a near-diploid dataset from Simulation A. For each tumor clone, the original and enhanced LRC of a randomly selected cell belonging to the clone are compared. The results suggest LRC data are significantly improved by the CAE model, and copy number information is accurately disentangled from technically confounded factors.

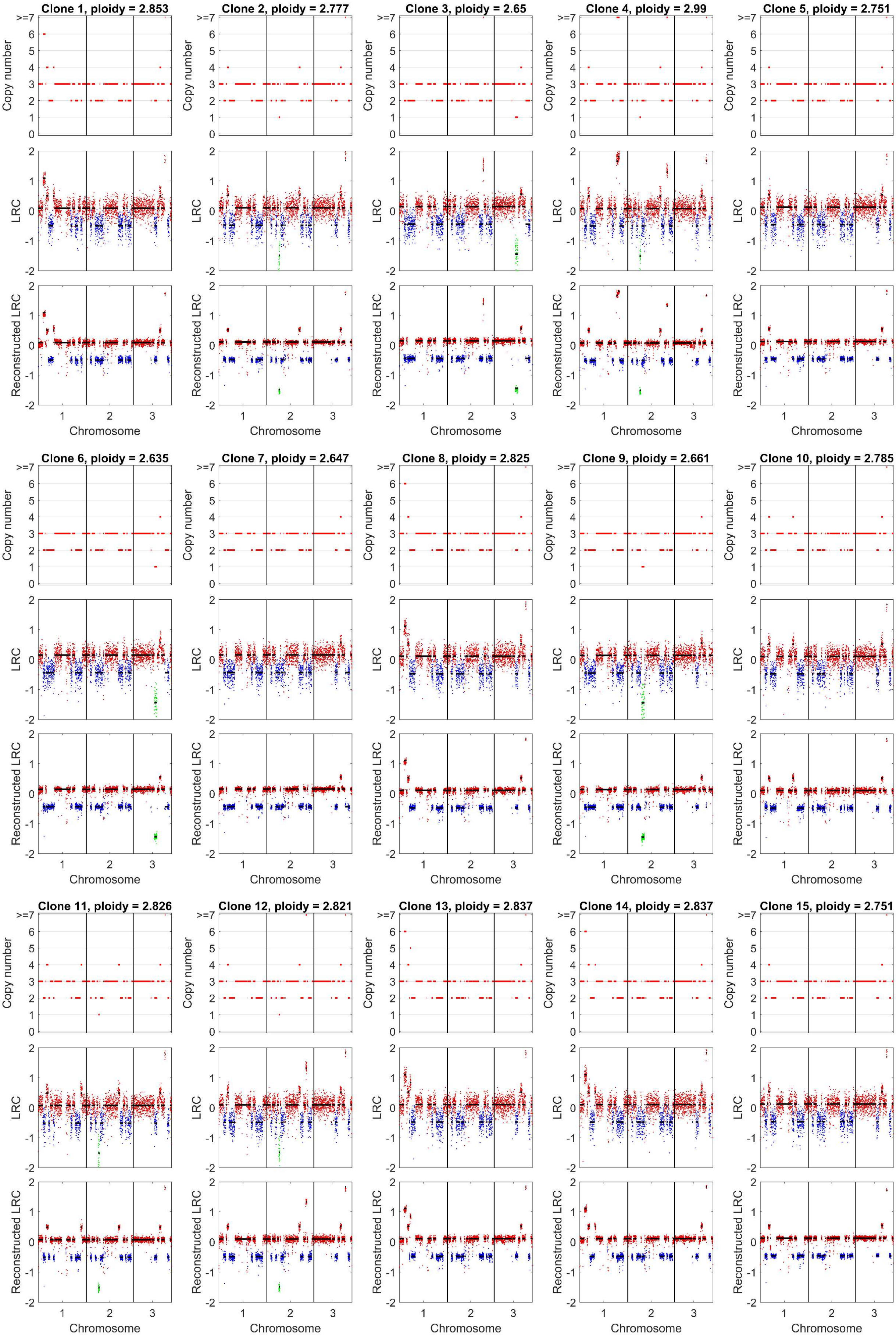

**Figure S3.** Copy number estimation results on a near-triploid dataset from Simulation B. For each tumor clone, the original and enhanced LRC of a randomly selected cell belonging to the clone are compared. The results suggest LRC data are significantly improved by the CAE model, and copy number information is accurately disentangled from technically confounded factors.

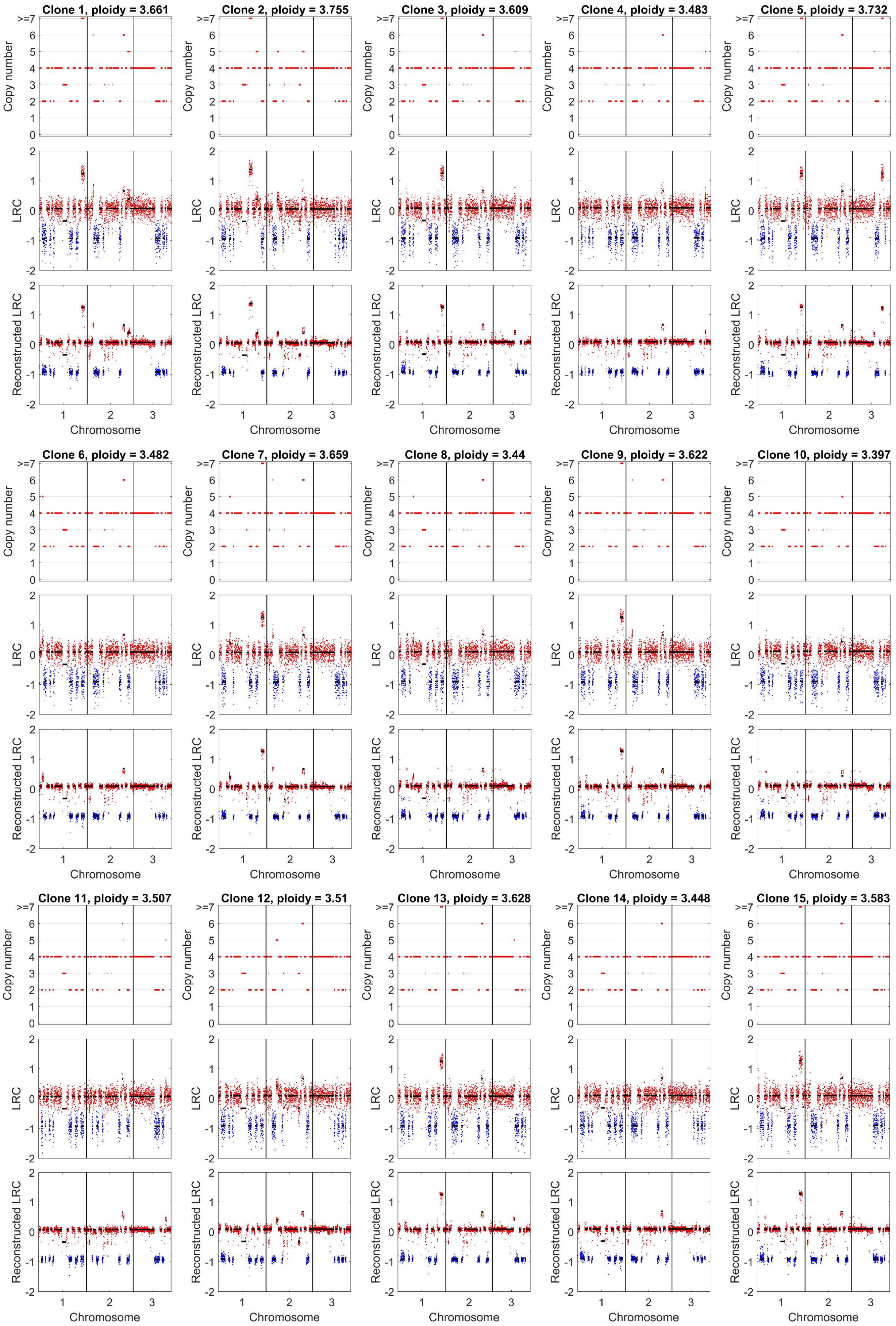

**Figure S4.** Copy number estimation results on a near-tetraploid dataset from Simulation C. For each tumor clone, the original and enhanced LRC of a randomly selected cell belonging to the clone are compared. The results suggest LRC data are significantly improved by the CAE model, and copy number information is accurately disentangled from technically confounded factors.

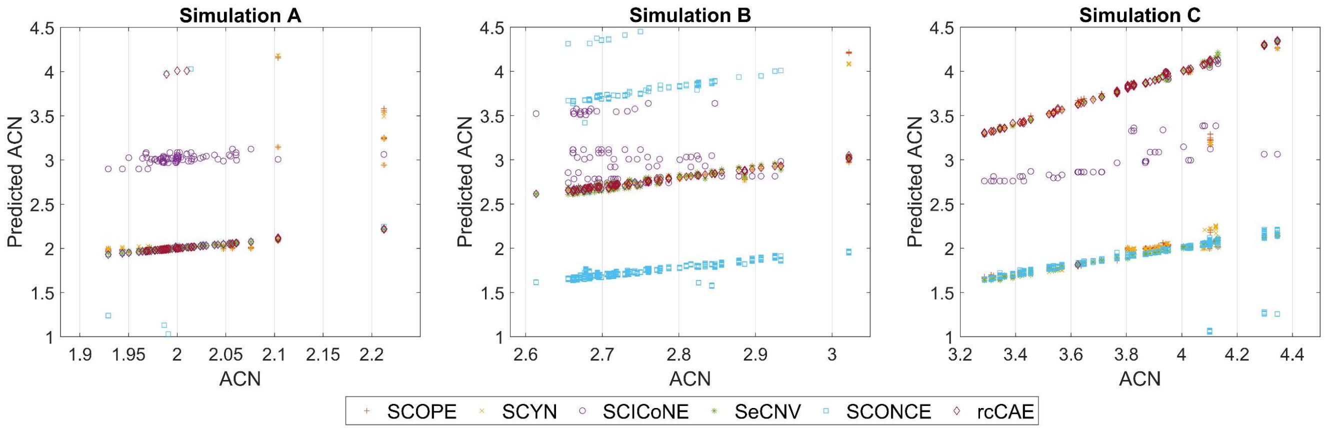

**Figure S5.** Comparison between the predicted and ground truth average copy numbers (ACNs). Results on simulated near-diploid datasets (Simulation A), near-triploid datasets (Simulation B) and near-tetraploid datasets (Simulation C) are analyzed. rcCAE yields better predictions than other methods across different simulations, especially on near-tetraploid datasets (Simulation C).

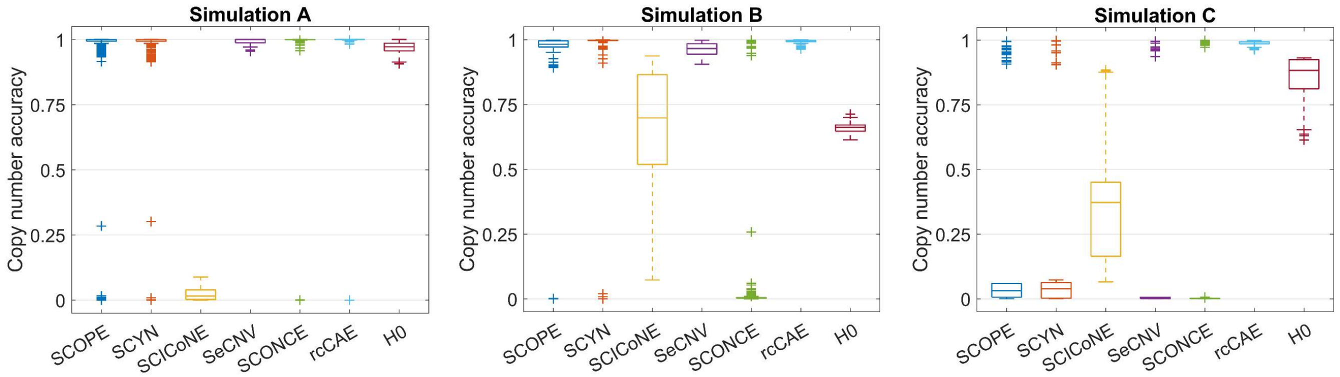

**Figure S6.** Copy number accuracy of the investigated methods in estimating absolute copy numbers. For each cell, copy number accuracy is calculated as the proportion of bins whose copy numbers are correctly predicted. Results on simulated 5 near-diploid datasets (Simulation A), 5 near-triploid datasets (Simulation B) and 5 near-tetraploid datasets (Simulation C) are analyzed. Each boxplot is generated based on all cells included in the corresponding simulation. The average number of bins used for evaluation is 2509 for Simulation A, 2636 for Simulation B and 2166 for Simulation C. H0 denotes the baseline model that predicts the most common copy number (2 for diploid, 3 for triploid and 4 for tetraploid datasets) for all bins.

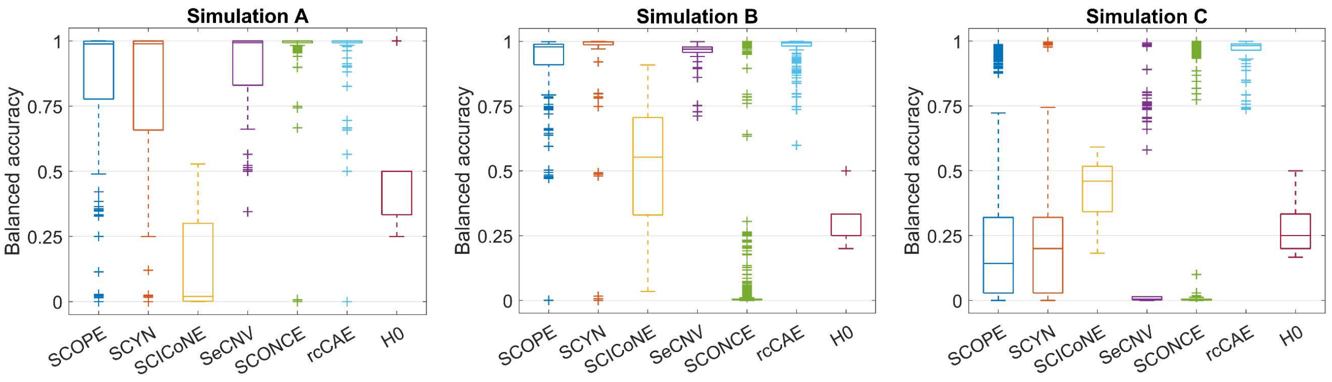

**Figure S7.** Balanced accuracy of the investigated methods in estimating absolute copy numbers. The balanced accuracy is defined as the mean of per-class accuracies. Results on simulated 5 near-diploid datasets (Simulation A), 5 near-triploid datasets (Simulation B) and 5 near-tetraploid datasets (Simulation C) are analyzed. Each boxplot is generated based on all cells included in the corresponding simulation. The average number of bins used for evaluation is 2509 for Simulation A, 2636 for Simulation B and 2166 for Simulation C. H0 denotes the baseline model that predicts the most common copy number (2 for diploid, 3 for triploid and 4 for tetraploid datasets) for all bins.

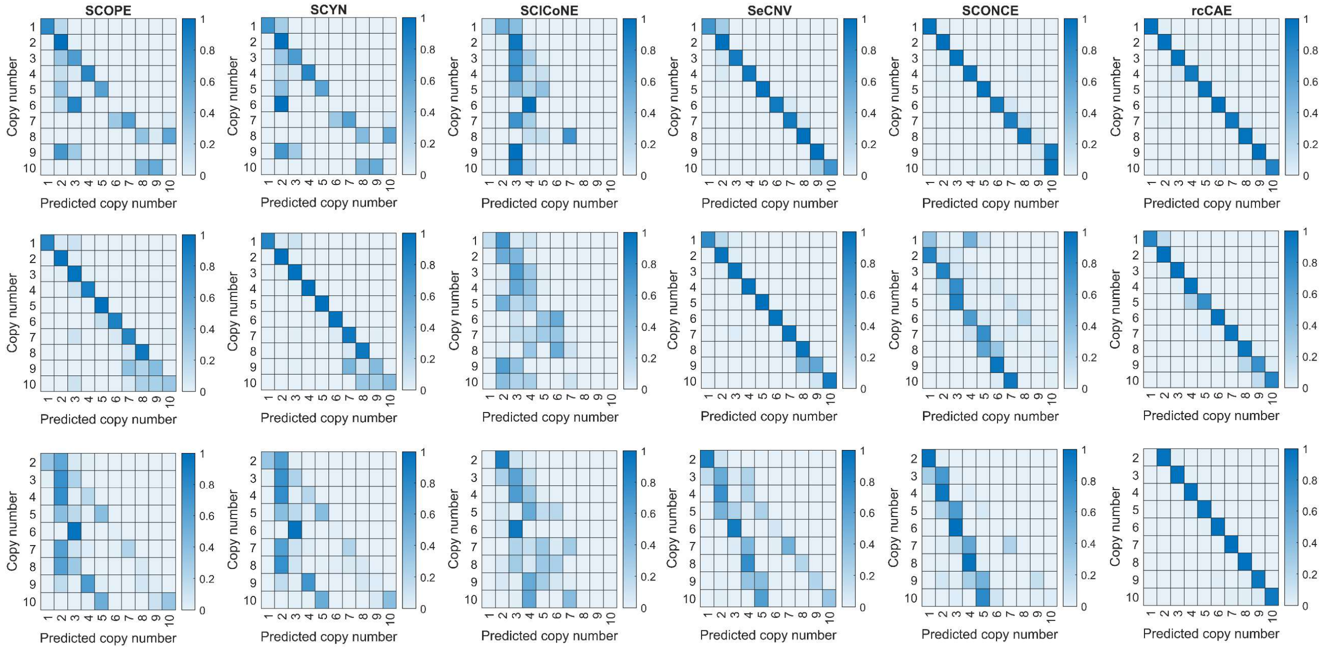

**Figure S8.** The accuracy of each method in identifying each copy number state. Results on simulated near-diploid datasets (top subfigures), near-triploid datasets (middle subfigures) and near-tetraploid datasets (bottom subfigures) are analyzed. It is observed that rcCAE makes accurate predictions for most cases especially on near-tetraploid datasets.

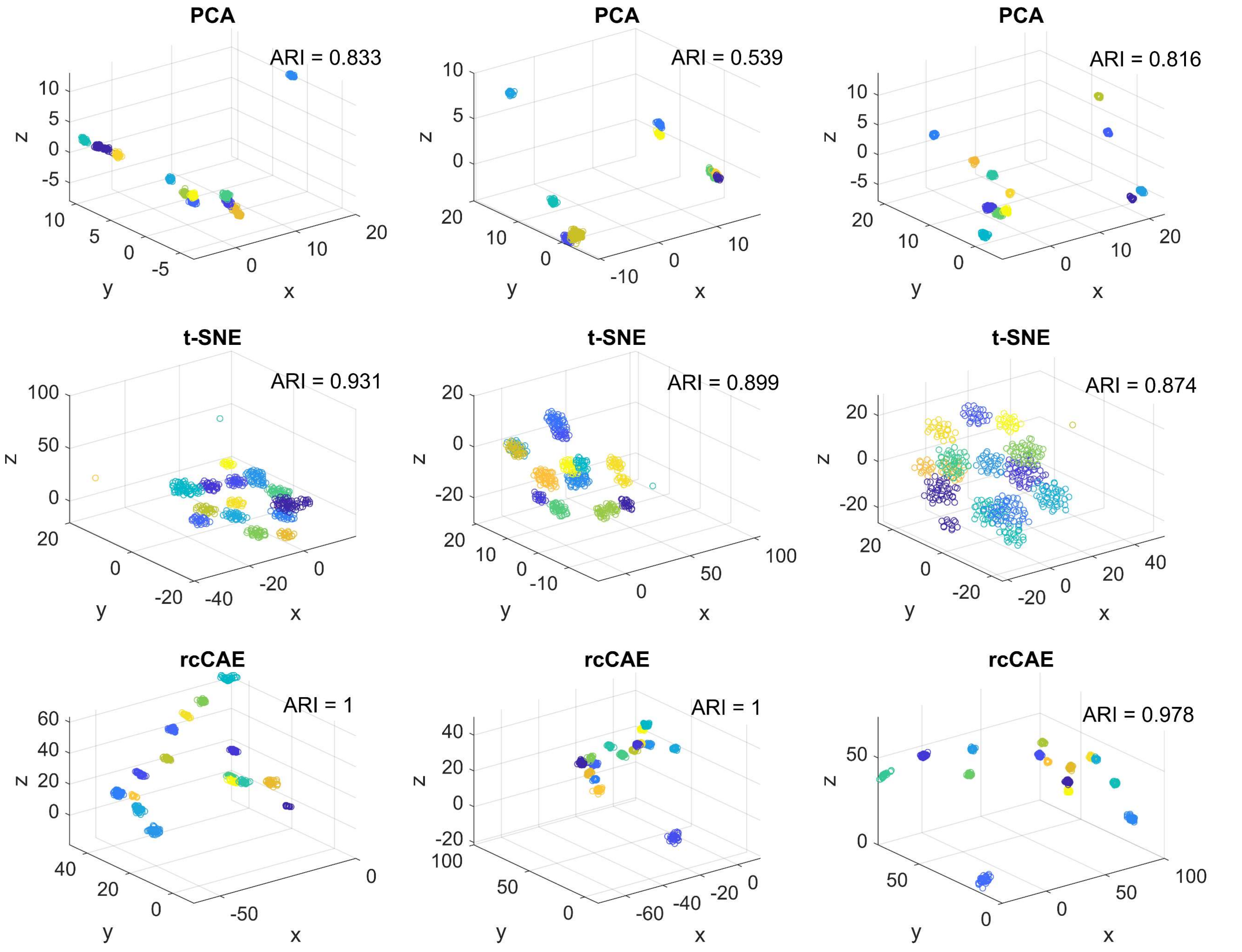

**Figure S9.** Clustering results of PCA, t-SNE and rcCAE on three simulated datasets. The left plots are results on a near-diploid dataset, the middle plots are results on a near-triploid dataset, and the right plots are results on a near-tetraploid dataset. Compared to PCA and t-SNE, rcCAE delivers more concentrated clusters of cells and yields higher ARI scores.

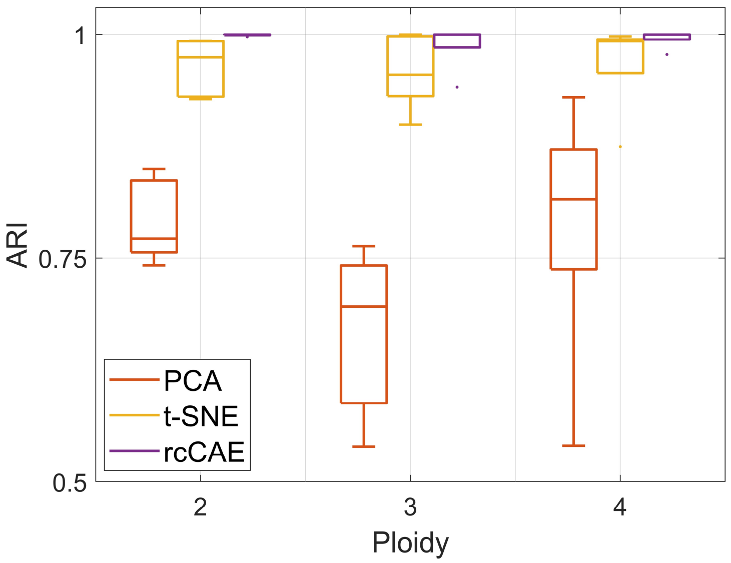

**Figure S10.** Comparison of clustering performance among PCA, t-SNE and rcCAE. Clustering of cells via a Gaussian mixture model is performed over the inferred 3-dimensional latent space for each of the methods. PCA is less effective in learning latent representations of cells, thus performs poorly in identifying clusters. rcCAE exactly recovers clonal architecture on 12 out of 15 datasets, and produces > 0.94 ARI scores on remaining 3 datasets.

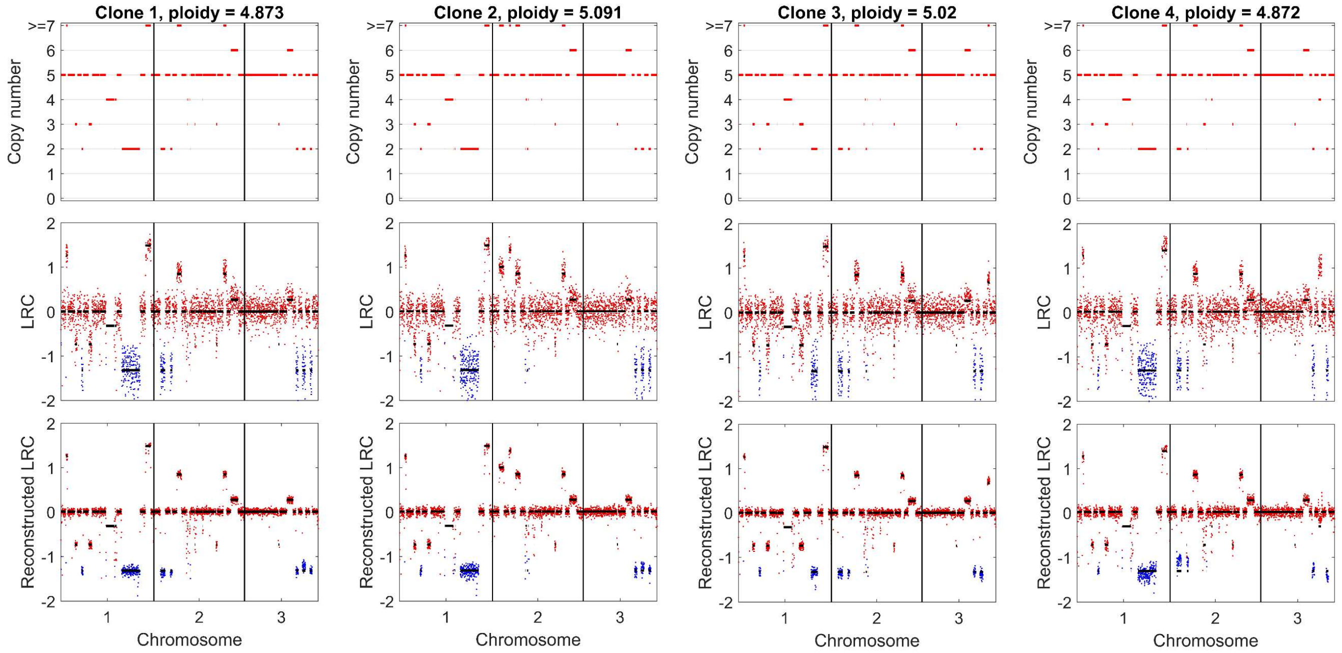

**Figure S11.** Copy number estimation results on a near-pentaploidy dataset from Simulation D. For each tumor clone, the original and enhanced LRC of a randomly selected cell belonging to the clone are compared. The results suggest LRC data are significantly improved by the CAE model, and copy number information is accurately disentangled from technically confounded factors.

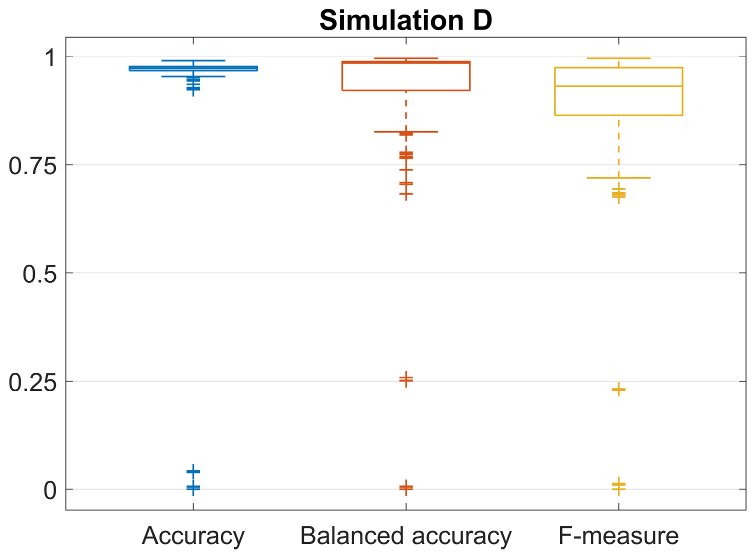

**Figure S12.** Copy number estimation performance of rcCAE on near-pentaploidy datasets (Simulation D). Each boxplot is generated based on all cells included in the simulation. The average number of bins used for evaluation is 2955.

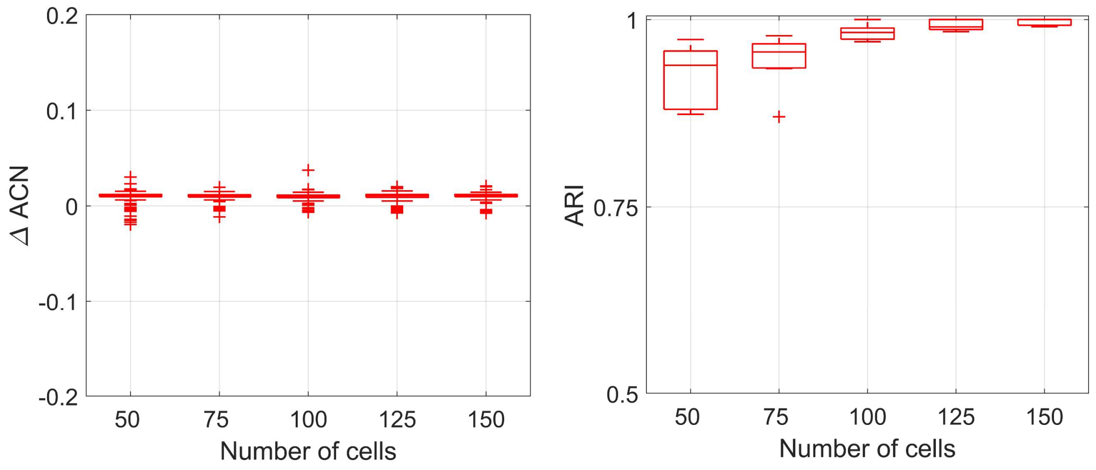

**Figure S13.** Ploidy estimation and cell clustering results of rcCAE on different-sized datasets. Values in {50, 75, 100, 125, 150} are tested for the number of cells, and 50 datasets for performance evaluation are generated from a near-diploid dataset in the Simulation A. Difference between the real and predicted ACNs (denoted as ΔACN) is calculated for each cell to indicate the ploidy estimation accuracy. Each boxplot is generated based on all cells included in the corresponding datasets.

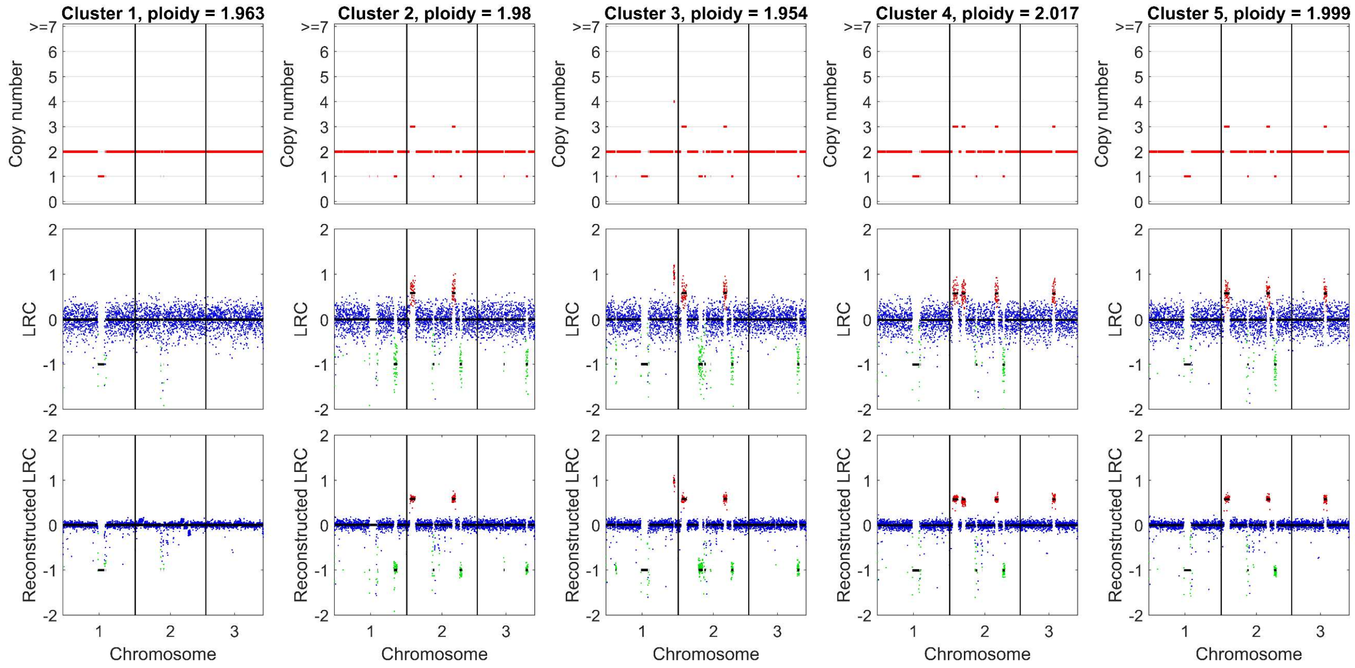

**Figure S14.** Copy number estimation results on a simulated dataset consisting of 75 cells. The results suggest rcCAE significantly enhances the quality of LRC data, and thus accurately estimates copy numbers for inferred segments.

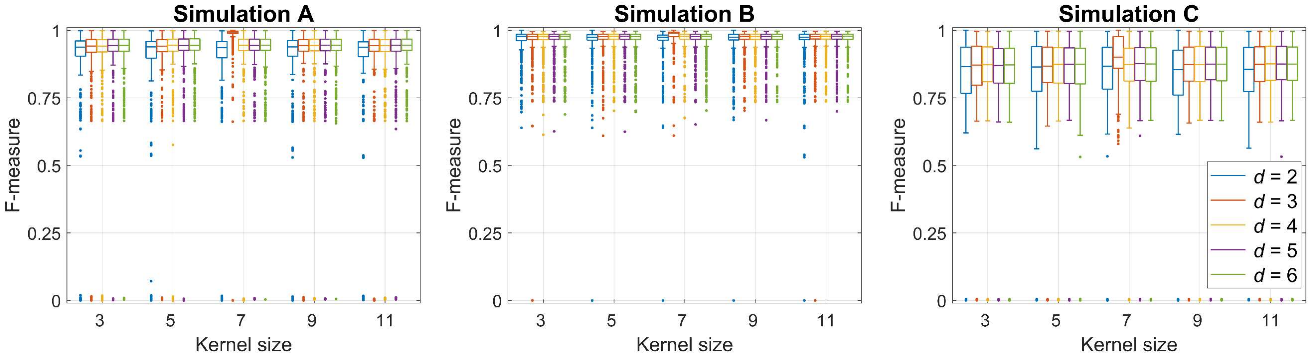

**Figure S15.** F-measure of rcCAE with respect to different configurations of kernel size and latent dimension. Values in {3, 5, 7, 9, 11} and {2, 3, 4, 5, 6} are tested for kernel size (denoted as *k*) and latent dimension (denoted as *d*), respectively. The results imply *k* and *d* have little effects on copy number estimation performance of rcCAE.

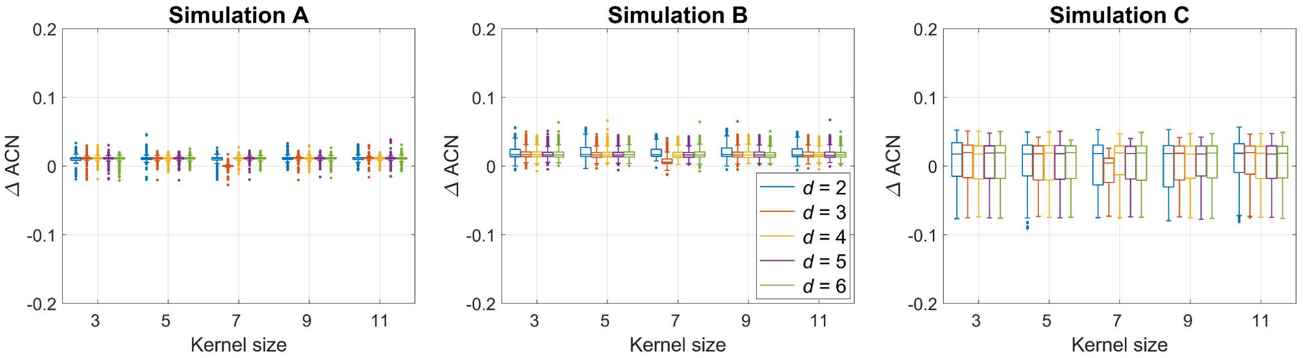

**Figure S16.** Ploidy estimation accuracy of rcCAE with respect to different configurations of kernel size and latent dimension. Values in {3, 5, 7, 9, 11} and {2, 3, 4, 5, 6} are tested for kernel size (denoted as *k*) and latent dimension (denoted as *d*), respectively. The results imply *k* and *d* have little effects on ploidy estimation accuracy of rcCAE.

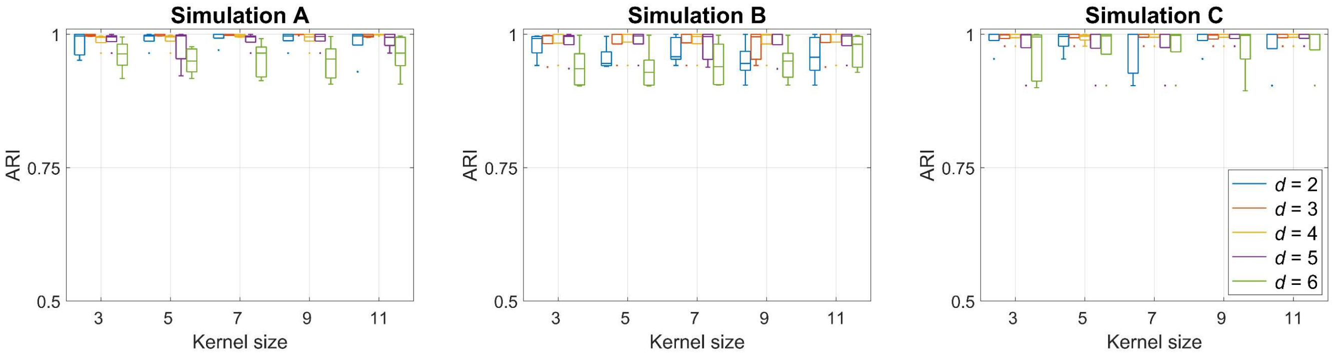

**Figure S17.** Clustering accuracy of rcCAE with respect to different configurations of kernel size and latent dimension. Values in {3, 5, 7, 9, 11} and {2, 3, 4, 5, 6} are tested for kernel size (denoted as *k*) and latent dimension (denoted as *d*), respectively. The results imply setting latent dimension to 3 or 4 delivers more accurate clustering results, and median-sized kernels tend to deliver more stable results.

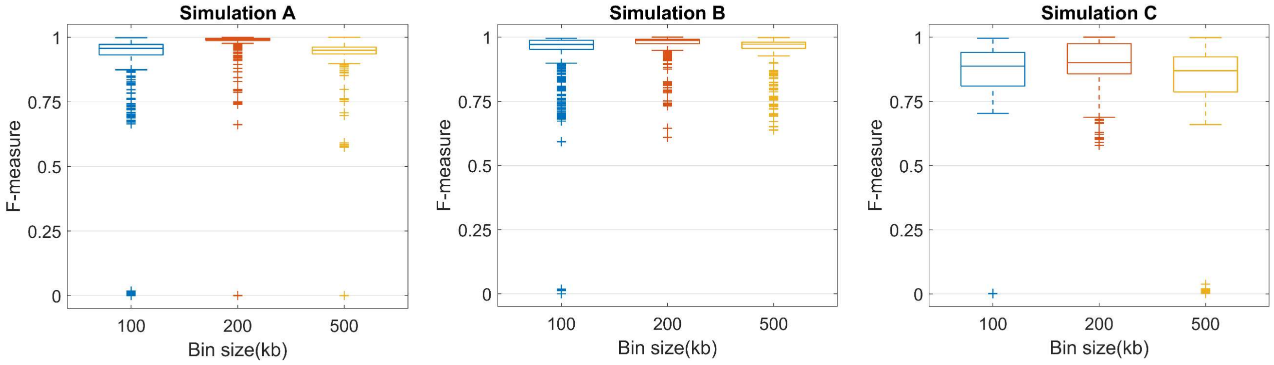

**Figure S18.** F-measure of rcCAE with respect to different values of bin size. Values in {100kb, 200kb, 500kb} are tested for the bin size. The results show bin size has no significant effect on copy number estimation performance of rcCAE.

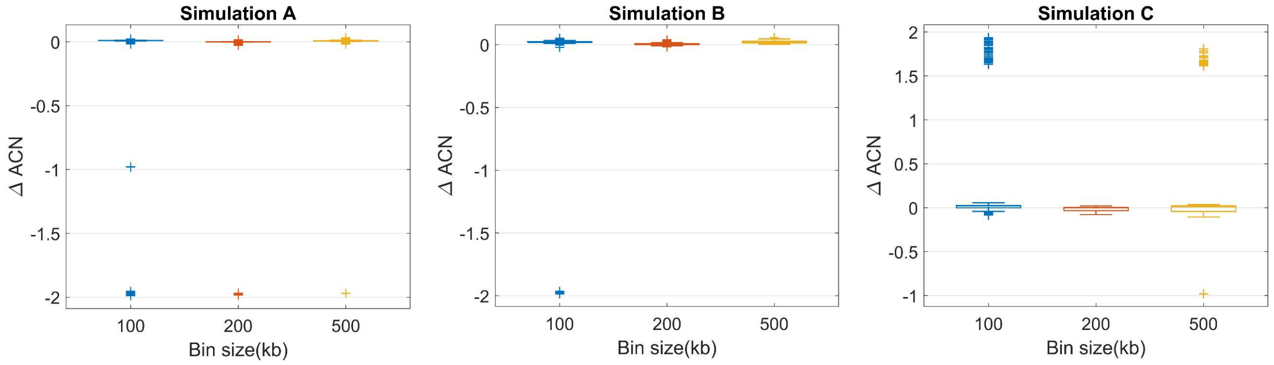

**Figure S19.** Ploidy estimation accuracy of rcCAE with respect to different values of bin size. Values in {100kb, 200kb, 500kb} are tested for the bin size. The results show bin size has little effect on ploidy estimation performance of rcCAE.

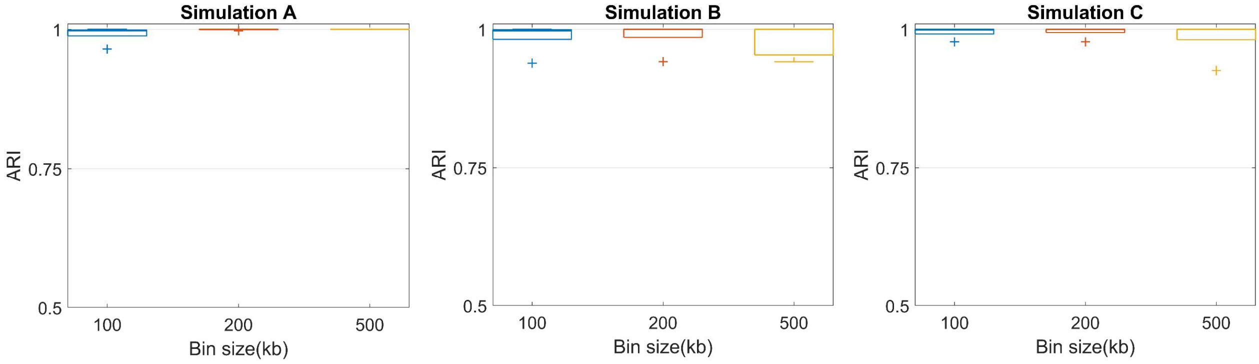

**Figure S20.** Clustering accuracy of rcCAE with respect to different values of bin size. Values in {100kb, 200kb, 500kb} are tested for the bin size. The results show bin size has little effect on clustering accuracy of rcCAE.

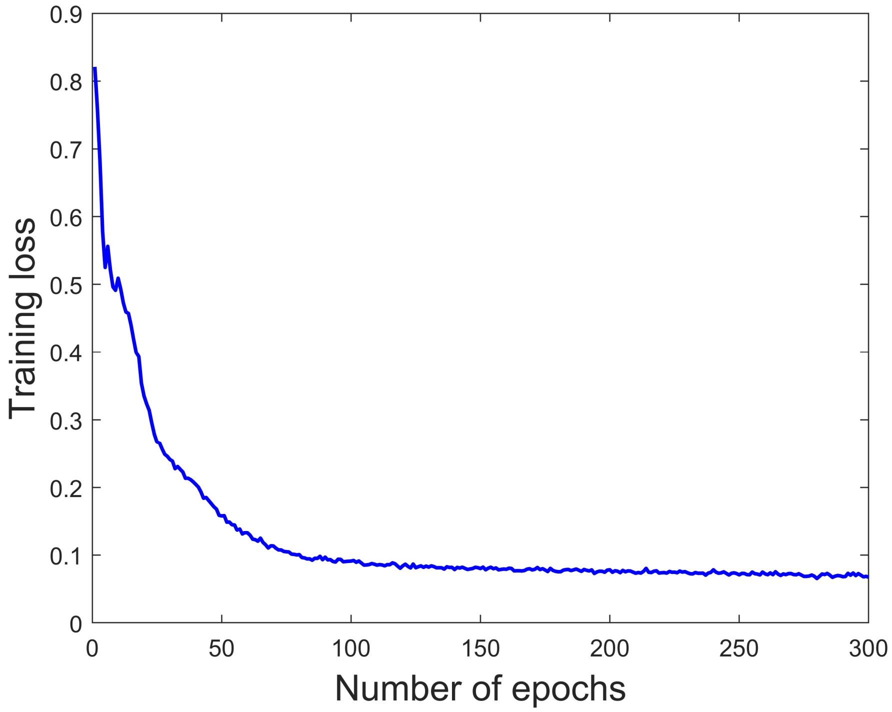

**Figure S21.** Training loss with respect to the number of epochs on breast cancer dataset. The training loss changes litter after 200 epochs, therefore the number of epochs for training the CAE is set to 200.

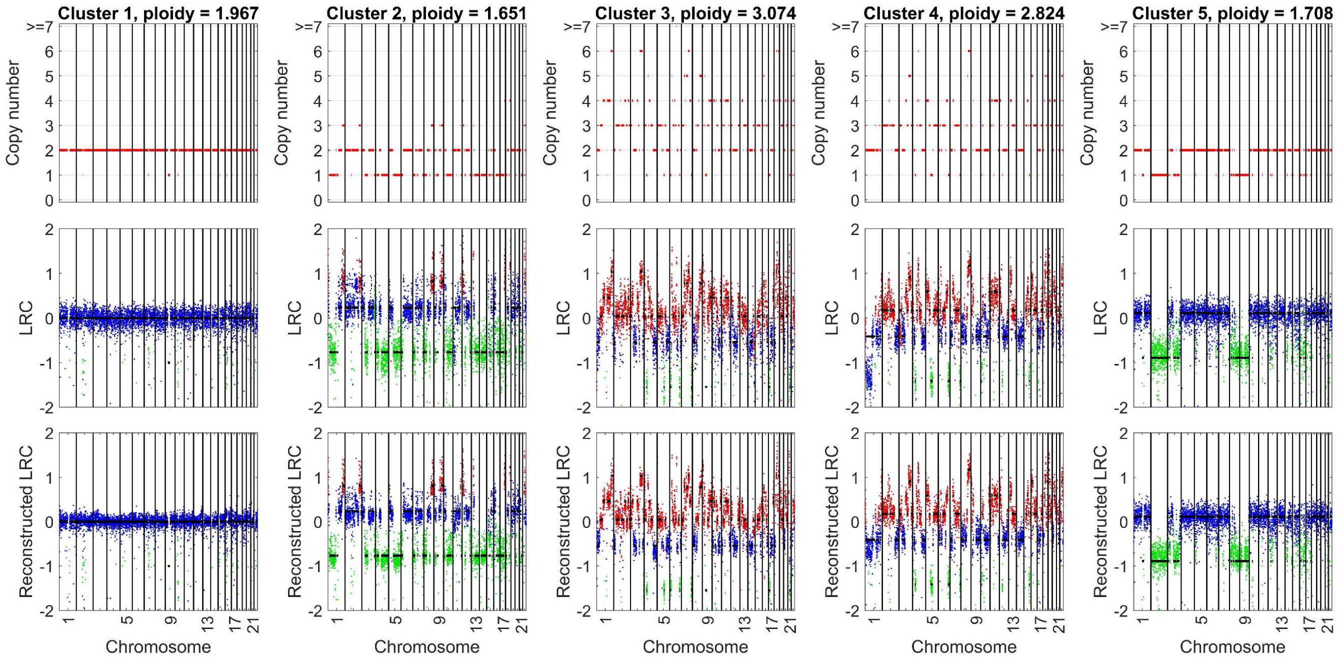

**Figure S22.** Copy number estimation results on breast cancer dataset. rcCAE identifies 5 clusters including one normal subpopulation and 4 tumor clones. Cluster 1 contains mainly diploid cells and represents the normal subpopulation. Cells from cluster 2 have hemizygous deletions on chromosomes 1p, 3p, 4q, 5q, 8p, 10q, 13q, 14, 16q and 17p. Cluster 3 consists of aneuploid cells that show copy number amplification on all chromosomes and hemizygous deletions on chromosomes 4-7. Cells from cluster 4 show copy number amplification on most of the chromosomes and hemizygous deletions on chromosomes 4-7. Only one cell is included in cluster 5, and this cell is significantly different from the cells of cluster 2 in copy number profiles, therefore is classified into a separate cluster by rcCAE.

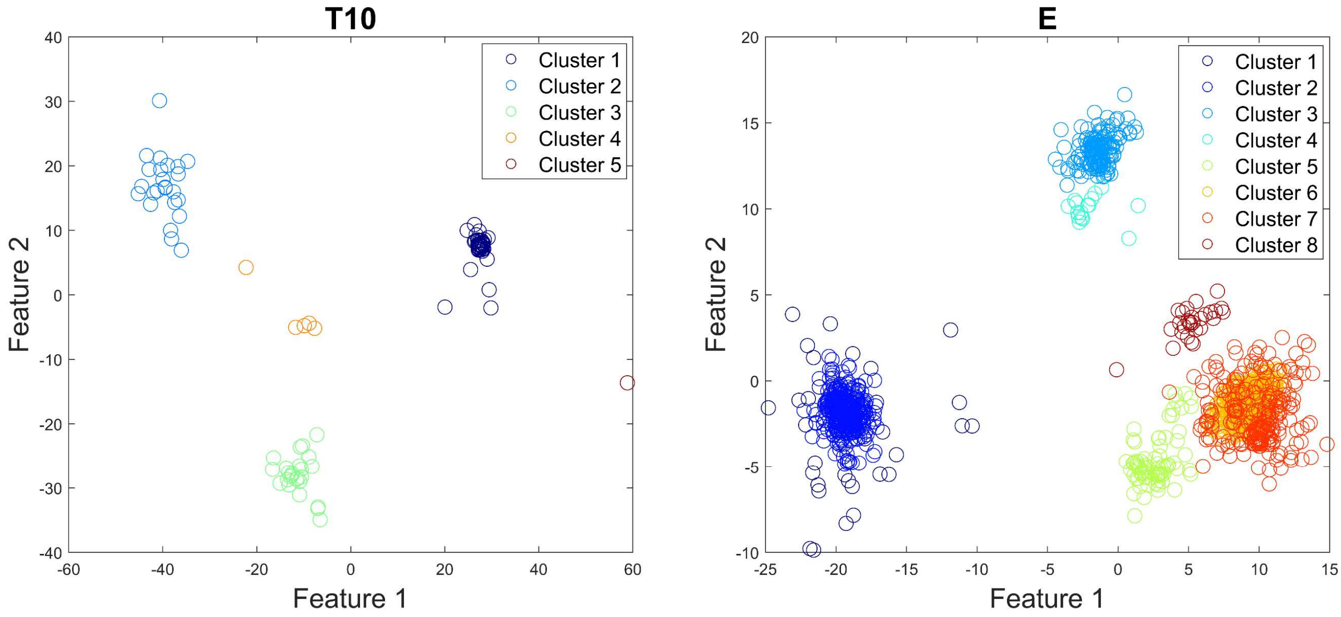

**Figure S23.** 2D visualization of the clustering results of rcCAE on real datasets. PCA is used to project the 3-dimensional latent data into a 2-dimensional space. For the 10X Genomics dataset (E), rcCAE generates over-partitioned results (clusters 1 and 2 may be from a same clone, clusters 3 and 4 are likely from a same clone, clusters 6 and 7 may come from a same clone).

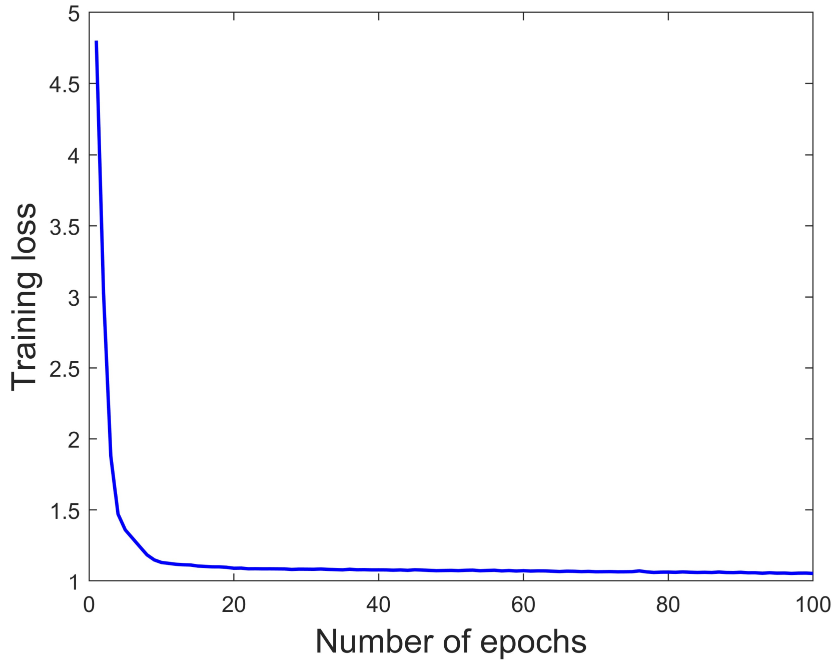

**Figure S24.** Training loss with respect to the number of epochs on 10X Genomics dataset. The training loss changes litter after 50 epochs, therefore the number of epochs for training the CAE is set to 50.
